# Time-Resolved Correspondences Between Deep Neural Network Layers and EEG Measurements in Object Processing

**DOI:** 10.1101/754523

**Authors:** Nathan C. L. Kong, Blair Kaneshiro, Daniel L. K. Yamins, Anthony M. Norcia

## Abstract

The ventral visual stream is known to be organized hierarchically, where early visual areas processing simplistic features feed into higher visual areas processing more complex features. Hierarchical convolutional neural networks (CNNs) were largely inspired by this type of brain organization and have been successfully used to model neural responses in different areas of the visual system. In this work, we aim to understand how an instance of these models corresponds to temporal dynamics of human object processing. Using representational similarity analysis (RSA) and various similarity metrics, we compare the model representations with two electroencephalography (EEG) data sets containing responses to a shared set of 72 images. We find that there is a hierarchical relationship between the depth of a layer and the time at which peak correlation with the brain response occurs for certain similarity metrics in both data sets. However, when comparing across layers in the neural network, the correlation onset time did not appear in a strictly hierarchical fashion. We present two additional methods that improve upon the achieved correlations by optimally weighting features from the CNN and show that depending on the similarity metric, deeper layers of the CNN provide a better correspondence than shallow layers to later time points in the EEG responses. However, we do not find that shallow layers provide better correspondences than those of deeper layers to early time points, an observation that violates the hierarchy and is in agreement with the finding from the onset-time analysis. This work makes a first comparison of various response features—including multiple similarity metrics and data sets—with respect to a neural network.

## 1. Introduction

In order to fully understand visual processing, we must be able to concretely model the computations that are carried out by the brain on not only synthetic but also natural stimuli. To this end, we can assess how well a proposed computational model of the visual system models actual neural computations by comparing it with neural data. There is much current interest in purely feed-forward convolutional neural networks (CNNs) as models to describe visual processing in humans and non-human primates (Cadena et al., 2019; Cichy et al., 2016; Yamins et al., 2014). These models are compelling because they were originally inspired by biological principles (Fukushima, 1980), and they are high performing in terms of the behaviour or task they were trained to perform. Furthermore, it has been shown that artificial neurons in these models correspond well to single-unit neural recordings (Bashivan et al., 2019; Cadena et al., 2019; Yamins et al., 2014; Yamins and DiCarlo, 2016).

The computations carried out in these kinds of models are hierarchical in nature in the sense that relatively simple features are encoded in the shallow layers and relatively complex features are encoded in the deeper layers. In particular, filters in a shallow layer of CNNs have Gabor-like receptive field properties that are similar to those that can be found in early visual cortex (Hubel and Wiesel, 1959; Marĉelja, 1980; Zeiler and Fergus, 2014). In addition, object-like features maximally activate artificial neurons in deeper layers of a CNN, implying that more complex features are extracted in those layers (Zeiler and Fergus, 2014).

In human neuroimaging studies, recent research has shown that features represented in the shallow layers of a CNN map to fMRI responses in early visual cortex, and features represented in the deeper layers map to responses from higher visual cortex (Güçlü and van Gerven, 2015; Seibert et al., 2016; Wen et al., 2017). Moreover, it has also been shown that stimulus representations in CNNs can be mapped temporally, where peak correspondence between a neural network and MEG responses occurred at earlier time points for shallow layers and at later time points for deeper layers (Cichy et al., 2016; Seeliger et al., 2017).

Representational similarity analysis (RSA) has emerged as a powerful method for comparing models to response data because it allows data collected from such different modalities as fMRI, MEG/EEG and single-unit recordings to be compared. This analysis framework has been used in a variety of contexts successfully (Cichy et al., 2016, 2017; Kaneshiro et al., 2015b; Khaligh-Razavi and Kriegeskorte, 2014; Kriegeskorte et al., 2008a; Yamins et al., 2014). When considering neural data, RSA operates on responses to *N* stimuli. Dissimilarity values are computed from responses to each pair of stimuli and summarized in a symmetric distance matrix termed a Representational Dissimilarity Matrix (RDM). The similarity metric, by which dissimilarities between pairs of stimuli are computed, is a key ingredient of RSA. However, its effects have not been studied extensively.

In this work, we compare EEG RDMs computed using three different similarity metrics with model RDMs and show that correlations between the EEG data and the neural network depend on the metric used. We focus on EEG, as it provides the temporal resolution to study visual processing dynamics, whereas fMRI does not allow us to record the fine-grained temporal details of a task. Furthermore, by investigating the model-EEG RDM correlations over the time course of the neural response, we find that they behave in a fashion that would not be expected in a purely feed-forward network: the onset of correlations associated with each layer are simultaneous rather than sequential, and shallow and deeper layers of the CNN correspond to RDMs computed from early time points of the EEG responses similarly well. This observation holds over independent data sets collected from two different laboratories using the same image set and response modality (Kaneshiro et al., 2015a; Cichy and Pantazis, 2017).

Our contributions are three-fold. Firstly, we study how representations obtained from EEG data compare with representations in the CNN, their implications on the model class in general and on the stimuli used in object processing experiments. Secondly, we show that derived correlation time courses depend on the similarity metric used to compute RDMs for RSA. Finally, we show that the high-level observations generalize across EEG data collected in different laboratories.

## 2. Methods

### 2.1. EEG Data Sets

We analyzed two publicly available EEG data sets containing responses to a common stimulus set of 72 images (shown in Figure 1). This image set is a subset of a larger 92-image set which has been used in several previous MEG and fMRI studies (Cichy and Pantazis, 2017; Khaligh-Razavi and Kriegeskorte, 2014; Kriegeskorte et al., 2008a,b), thus enabling broad comparison of results. In the 72-image set, each image belongs to one of six categories (12 images per category): human body (HB), human face (HF), animal body (AB), animal face (AF), fruits and vegetables (FV) and inanimate objects (IO).

**Figure 1:**
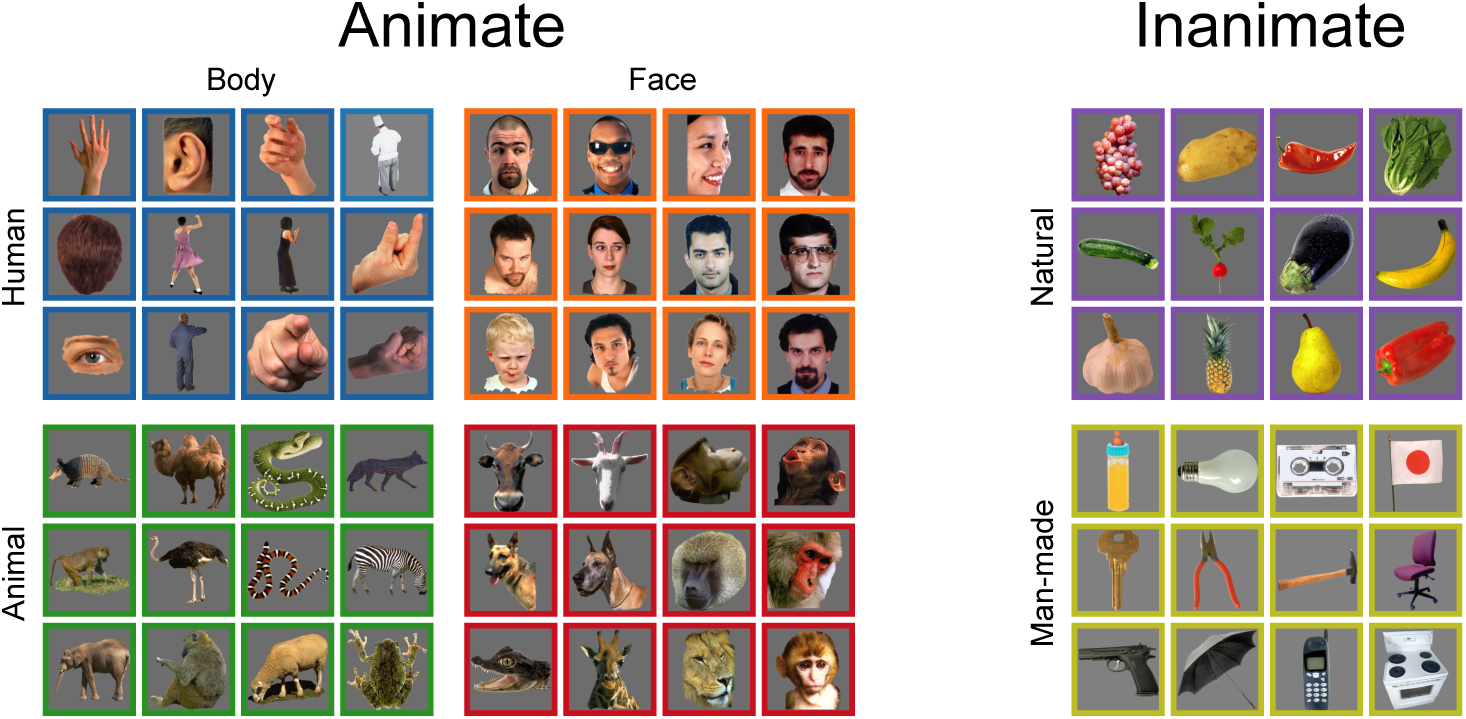
EEG stimulus set of 72 images. Both EEG data sets include responses to a shared stimulus set, whose images fall under six animate and inanimate categories. Coloured borders are included to illustrate category membership only, and were not presented during data collection. Adapted from Kaneshiro et al. (2015b).

The first data set (Kaneshiro et al., 2015a) has been previously analyzed in a published study on object processing (Kaneshiro et al., 2015b). This data set will henceforth be referred to as “Data Set 1”. Briefly, the data were collected from 10 subjects using a 128-electrode array. Each subject passively viewed each of the 72 stimuli 72 times (*N*_trials_ = 72). Each stimulus spanned a visual angle of 7*^◦^ ×* 6.5*^◦^* and was shown for 500 ms followed by a 750-ms interval between the presentation of each stimulus. For a more detailed description of the data acquisition process, see Kaneshiro et al. (2015b).

The second data set was analyzed in a previous MEG/EEG study (Cichy and Pantazis, 2017). This data set—henceforth referred to as “Data Set 2”—was included so that the robustness of our results could be assessed across laboratories. This data set comprises 74-electrode responses from 16 subjects viewing each of 92 stimuli (full set) 28 times (*N*_trials_ = 28). Our analyses consider responses to only the 72 images common to both stimulus sets. Each stimulus spanned a visual angle of 4*^◦^* and was shown for 500 ms followed by an interval of 1000 or 1100 ms before the presentation of the next stimulus. More information on data acquisition is provided in Cichy and Pantazis (2017).

### 2.2. EEG Preprocessing and Noise Normalization

We applied the same preprocessing and noise normalization procedures to the individual recordings of both EEG data sets as follows. Firstly, each recording was high-pass filtered at 0.3 Hz, low-pass filtered at 25 Hz, and temporally downsampled by a factor of 16, reducing the sampling rate from 1 kHz at acquisition to 62.5 Hz (16 ms per time sample). Ocular artifacts were removed using a regression procedure, and electrodes on the face were excluded from further analysis in Data Set 1, reducing the channel dimension for these recordings from 128 to 124. Data were epoched into trials from *−*112–512 ms relative to stimulus onset. Large noise transients were replaced with the spatial average of data from neighboring electrodes. Finally, data were converted to average reference, and every trial of data was baseline corrected on a per-electrode basis based on mean activity between *−*112–64 ms relative to stimulus onset.

Following preprocessing, we converted single-trial responses to pseudo-trials in order to improve the SNR of the data (Guggenmos et al., 2018). Five pseudo-trials were computed for each stimulus, using random partitioning of the trials; averaging was performed on a per-subject, per-stimulus basis. Inherent to EEG is the fact that each electrode has different levels of noise due to, for instance, an electrode’s location on the scalp. Moreover, correlations may exist among responses recorded from different electrodes. Thus, noise normalization is recommended as a general preprocessing step in order to reduce these effects on downstream analyses (Guggenmos et al., 2018). We performed multivariate noise normalization to whiten the data, so that electrodes that were relatively less noisy would be emphasized over electrodes that were relatively more noisy. Similarly to Guggenmos et al. (2018), we computed the sample covariance matrices using the “epoch” method, whereby covariance matrices were computed at each time point and then averaged across time.

After preprocessing, the dimensions of Data Set 1 were 10 subjects *×* 72 stimuli *×* 72 trials *×* 40 time points *×* 124 electrodes, and the dimensions of Data Set 2 were 16 subjects *×* 72 stimuli *×* 28 trials *×* 40 time points *×* 74 electrodes.

### 2.3. Test-Retest Reliability of Neural Data

The reliability of the EEG data imposes an upper limit on how well any model can explain the data. Therefore, before performing any downstream analyses, we assessed the quality of the EEG data by computing the test-retest reliability for each electrode at each time point. On a per-subject basis, we computed the reliability over 10 random permutations. In each permutation, we randomly split the *N*_trials_ trials in half, resulting in two partitions with *N*_trials_*/*2 trials each. Therefore, for each electrode and for each time point, the dimensions of the data were 72 stimuli *× N*_trials_*/*2 trials. For each of the two data partitions (each with half the total number of trials), we computed the mean across the trials, resulting in two vectors of length 72. The uncorrected reliability, *R*, was then the correlation coefficient between the two vectors for a particular electrode. Spearman-Brown correction was applied to this value using the following formula:

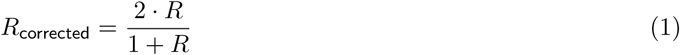

The reliability at each time bin was defined to be the average reliability across all electrodes. For each subject, the mean of the corrected reliability, *R*_corrected_, was computed across all permutations. Finally, the mean and standard error of the mean were computed across the subject pool separately for each data set.

In order to compare the reliability of the two data sets, we controlled for the number of trials used in the reliability analysis. By combining the trials across subjects, we computed the reliability of the data sets created using subsets of all the trials, which we call “pseudo-data sets”. This allowed us to observe how reliability of the pseudo-data set changes with increasing number of trials at a particular time point in the EEG time series, which we chose to be 144 ms, since this is the time point where the time-varying reliability is approximately maximal. Here the reliability is also defined to be the average reliability across all electrodes. This procedure was repeated 100 times, where each repetition corresponded to a data set curated from randomly chosen trials without replacement. Finally, the mean and standard deviation across the 100 randomly selected pseudo-data sets were computed.

### 2.4. Representational Similarity Analysis

Representational similarity analysis (RSA) is a framework through which data from different modalities can be compared (Kriegeskorte et al., 2008a). Cross-modal comparisons are enabled by abstracting modality-specific responses to representational dissimilarity matrices (RDMs), which summarize dissimilarities among all pairs of stimuli. Guggenmos et al. (2018) have shown that an important ingredient for RSA is the similarity metric used to compute the pairwise dissimilarities for the RDM. We investigated three commonly used metrics: pairwise decoding (classification) accuracy, cross-validated Euclidean distance and cross-validated Pearson correlation. Our computations of these similarity metrics were modelled after code provided in Guggenmos et al. (2018). As described extensively in that study, for each of these cross-validated similarity measures, one randomly selected pseudo-trial of each stimulus class was left out for validation when computing the pairwise dissimilarity value. This procedure was repeated 20 times with randomly computed pseudo-trials in each iteration, resulting in 20 dissimilarity values computed for each pair of stimuli. We report the mean dissimilarities across the 20 iterations. In order to obtain time-resolved RDMs for the EEG data, we computed each of the three similarity metrics at every time sample of the response data, resulting in 40 temporally resolved RDMs for each EEG data set and similarity metric for each subject.

#### 2.4.1. Decoding Accuracy

When pairwise decoding accuracy was used as a similarity metric, a classifier was trained to discriminate between EEG responses to each pair of images (Cichy et al., 2016; Guggenmos et al., 2018). Here we trained a linear discriminant analysis classifier (LDA) to classify responses to all pairs of stimuli. A poor classification accuracy for a given pair (near two-class chance level of 0.5) suggests relatively higher similarity in the neural response representation space; on the other hand, a high classification accuracy closer to 1.0 would indicate relatively less similarity in the neural response representation space. The RDM was constructed from the set of pairwise accuracies, with 0.5 subtracted from each classification accuracy so that RDM values ranged from 0 to 0.5.

#### 2.4.2. Cross-validated Euclidean Distance

Cross-validated squared Euclidean distance was introduced in Guggenmos et al. (2018) as a metric for assessing similarity between pairs of neural responses in the native units of squared voltage differences. Note that when the cross-validated Euclidean distance metric is paired with multivariate noise normalization, the metric is equivalent to the cross-validated Mahalanobis distance (crossnobis distance, Walther et al. (2016)). The equation is defined as follows:

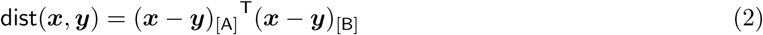

where [A] and [B] represent two parts of the data partitioned on the pseudo-trial dimension and where ***x*** and ***y*** represent the neural response vectors, each to a particular stimulus.

#### 2.4.3. Cross-validated Pearson Correlation

The cross-validated Pearson correlation was introduced for EEG and MEG applications in Guggenmos et al. (2018) as a unit-free similarity metric between neural responses. This analysis was carried out on the pseudo-trial data. The equation is defined as follows:

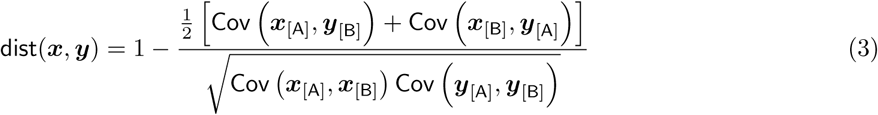

As before, [A] and [B] represent two partitions of the data and ***x*** and ***y*** represent the neural response vectors.

### 2.5. Computational Model

As a computational model, we used VGG19 (Simonyan and Zisserman, 2014), which was previously trained on the ImageNet data set (Deng et al., 2009). It is one of the best-performing neural networks for object categorization (He et al., 2016). This network consists of five blocks of convolutions and pooling and three fully connected layers. In each of the blocks, there are either two or four convolutional layers followed by a max pooling stage. The features from five pooling layers (`pool1, pool2, pool3, pool4, pool5`) and the penultimate fully connected layer (`fc2`) were selected from the model for comparison with the EEG data.

We computed RDMs for the representative layers of the CNN using Pearson correlation as the similarity metric (Kriegeskorte, 2009; Yamins et al., 2014). At each representative layer, the feature activation map for a particular image was vectorized and then correlated with the feature activation maps obtained from every other image.

### 2.6. Comparing the Model with Neural Data Using RSA

At each time bin of the EEG response, the RDMs computed from the model were compared to the RDMs from the EEG data by computing Spearman’s correlation between the upper triangular portions of the EEG and model RDMs. Doing this across all time bins resulted in a correlation time course.

### 2.7. Computing the Noise Ceiling of the Correlation Time Course

The noise ceiling of the correlation time course is a measure computed using the EEG data, representing the maximum attainable correlation that the data RDM at each time bin can have with any other RDM. Here we used a method described extensively in Nili et al. (2014) to compute the noise ceiling. Briefly, the following steps were performed at each time bin. First, the upper triangular portion of each subject’s EEG RDM was rank transformed and vectorized. We used a leave-one-subject-out approach: using the upper triangular rank transformed data vector of all the subjects’ EEG RDMs, each subject’s data vector was Spearman rank correlated with the average data vector of all other subjects resulting in *N*_subjects_ correlation values. The noise ceiling at the particular time bin was then computed by averaging the rank correlation across subjects.

### 2.8. Computing Time of Correlation Onset and Peak

In this section, we describe how the correlation onset time between model RDMs and EEG RDMs was determined. For each correlation time course, a two line segment piece-wise linear function was fit for the correlation values starting at *−*112 ms and ending at approximately the time point at which the correlation time course reaches peak correlation (as in Kohler et al. (2016)). For Data Set 1, this end point was at 112 ms, and for Data Set 2, it was at 96 ms. The correlation onset time was then determined to be the time point at which the two segments of the piece-wise linear function intersected. The Python package described in Jekel and Venter (2019) was used to perform the piece-wise linear fit. The time of peak correlation is defined to be the time point at which the maximum correlation between the EEG and layer RDM occurs. We performed these computations across 1000 bootstrap samples, across subjects, and then computed the mean and error of the mean for each time bin. We report the linear regression *p*-value for each correlation metric (peak or onset), data set and similarity metric.

### 2.9. Optimization Procedures

EEG responses at each electrode could be the result of a mixture of neural activity that may or may not be relevant to the stimulus. Even if we assume that the mixing of the activity is linear, computing RDMs is problematic due to the fact that linear reweighting of neural responses could modify the resulting RDM almost arbitrarily. As will be shown in Section 3.3.1, this would result in relatively low correlations between the model and the neural data since the model (without any linear reweighting) is expected to explain neural responses that have been mixed. The two optimization procedures presented in the following sections use the data to account for this mixing matrix so that its effect on the model-to-data comparison is reduced. This approach was first applied by Khaligh-Razavi and Kriegeskorte (2014), where fMRI responses from humans and single-cell recordings from monkeys were used. The authors observed that using a linear re-scaling of features from several layers of a CNN resulted in RDMs that had higher correspondence to monkey and human IT RDMs than that of RDMs computed without linear re-scaling of features.

#### 2.9.1. Linearly Combining Features from the Neural Network

We sought to compute an optimal linear combination of features from a particular layer of the CNN such that the resulting layer RDM would be maximally correlated with the EEG RDM at a particular time point. We refer to this optimization procedure as the “linear combination optimization procedure”. Note that this is different from the method used in Khaligh-Razavi and Kriegeskorte (2014) since we learn a lower-dimensional set of features that are linear combinations of the original set of features. Concretely, at each time point *t*, and for each layer *l*, we performed the following optimization:

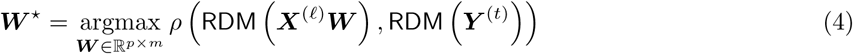

where *ρ*(***x, y***) is the Pearson correlation between ***x*** and ***y***, RDM(·) is the function that computes the vector of similarities between pairs of stimulus features from EEG or from the CNN, ***X***^(*l*)^*∈* R^72*×p*^, with *m < p*, are the vectorized stimulus features for layer *l*, ***W\**** is the optimal weight matrix and ***Y*** ^(*t*)^*∈* R^72*×k*^ are the EEG stimulus features at time *t*. Recall that 72 stimuli were used in this study.

Equation 4 was optimized using stochastic gradient descent (Bottou, 2010; Robbins and Monro, 1951) with a learning rate of *η* = 0.001 and a stopping criterion of *∈* = 0.001. This means that we stopped optimizing when the change in Pearson correlation computed from the previous iteration and from the current iteration of the optimization was less than or equal to 0.001. The optimization procedure was implemented using a Python package known as PyTorch (Paszke et al., 2017). The number of latent dimensions, *m*, was set to 1000 for all six layers used in the analyses. We performed four-fold cross-validation at each time point so that in each fold, the optimization was performed using 54 stimuli, resulting in an optimal weight matrix, and testing was performed on the remaining 18 stimuli not used in the optimization. In each fold, an RDM was computed with the test set stimuli using “new” stimulus features obtained by multiplying the optimal weight matrix with the original stimulus features (after this operation, the features for each stimulus were of length *m*). Using Spearman’s rank correlation, this “test set” RDM was then correlated to the “test set” EEG RDM from the particular time point that was being optimized for. The four-fold cross-validation procedure at each time point therefore resulted in four Spearman’s rank correlation values which were subsequently averaged. Note that the EEG RDMs were computed using three similarity metrics described previously and that the layer RDMs were computed using only the Pearson correlation similarity metric, as before.

#### 2.9.2. Scaling Features Element-Wise from the Neural Network

The method described in Section 2.9.1 is time consuming due to the number of weights that are being learned, which could be on the order of 10^8^. To reduce the computation time, we implemented an alternative method that is much faster. For each layer of the CNN, we sought to find a set of weights that *scaled* each element of the CNN feature vector. This is in contrast to the optimization problem described in Section 2.9.1 where we sought to find *linear combinations* of CNN features. The number of weights that were optimized in this method were at most on the order of 10^5^. We refer to this optimization procedure as the “element-wise scaling optimization procedure”. Concretely, at each time point *t*, and for each layer *l*, we performed the following optimization:

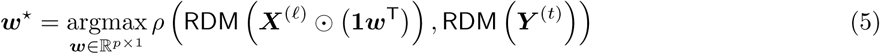

where ⊙ is the element-wise product operation, ***w\**** is the optimal scaling vector, **1** *∈* R^72*×*1^ is a column vector of ones, *p* is the dimension of the vectorized CNN features for a stimulus and as previously described in Section 2.9.1, ***X***^(*l*)^ *∈* R^72*×p*^ are the stimulus features for layer *l*, ***Y*** ^(*t*)^*∈* R^72*×k*^ are the EEG stimulus features at time *t*, *ρ*(***x, y***) is the Pearson correlation between ***x*** and ***y*** and RDM(·) is the function that computes the vector of similarities between pairs of stimulus features from EEG or from the CNN.

The optimization and cross-validation procedures for Equation 5 were the exact same as those described in Section 2.9.1, where the learning rate was set to *η* = 0.001, the stopping criterion was set to *E* = 0.001 and four-fold cross-validation was performed. This objective function and optimization procedure is similar to that implemented in Seibert et al. (2016).

## 3. Results

### 3.1. Neural Data Reliability

As noted previously in Section 2.3, test-retest reliability of the EEG data imposes an upper limit on how well an optimal model can explain the data. In our present analyses, we found that the test-retest reliability of the EEG data varies with time. Figure 2A shows the reliability of the neural responses across time for both data sets. In Data Set 1, we observed low reliability before the onset of the evoked response at approximately 80 ms, peak reliability in the time range of the evoked response (Luck and Kappenman, 2011) and a decrease in reliability gradually thereafter. The reliability was near maximal (approximately 0.8) between 100 and 200 ms. The reliability of Data Set 2 was also found to vary across time. Similarly to the observations from Data Set 1, we observed low reliability before the onset of the evoked response and maximal reliability (approximately 0.4) between 100 ms and 200 ms with a gradual decrease thereafter. Consistently across the two data sets, we observed that the reliability of the neural data was near maximal in the time range in which the evoked response occurs and near 0 around stimulus onset time. We note that Data Set 2 appears to be less reliable than Data Set 1 based on Figure 2A. This is most likely the result of there being fewer trials collected per stimulus in Data Set 2 as compared to that of Data Set 1 (72 trials vs. 28 trials per stimulus for Data Set 1 and Data Set 2 respectively).

**Figure 2:**
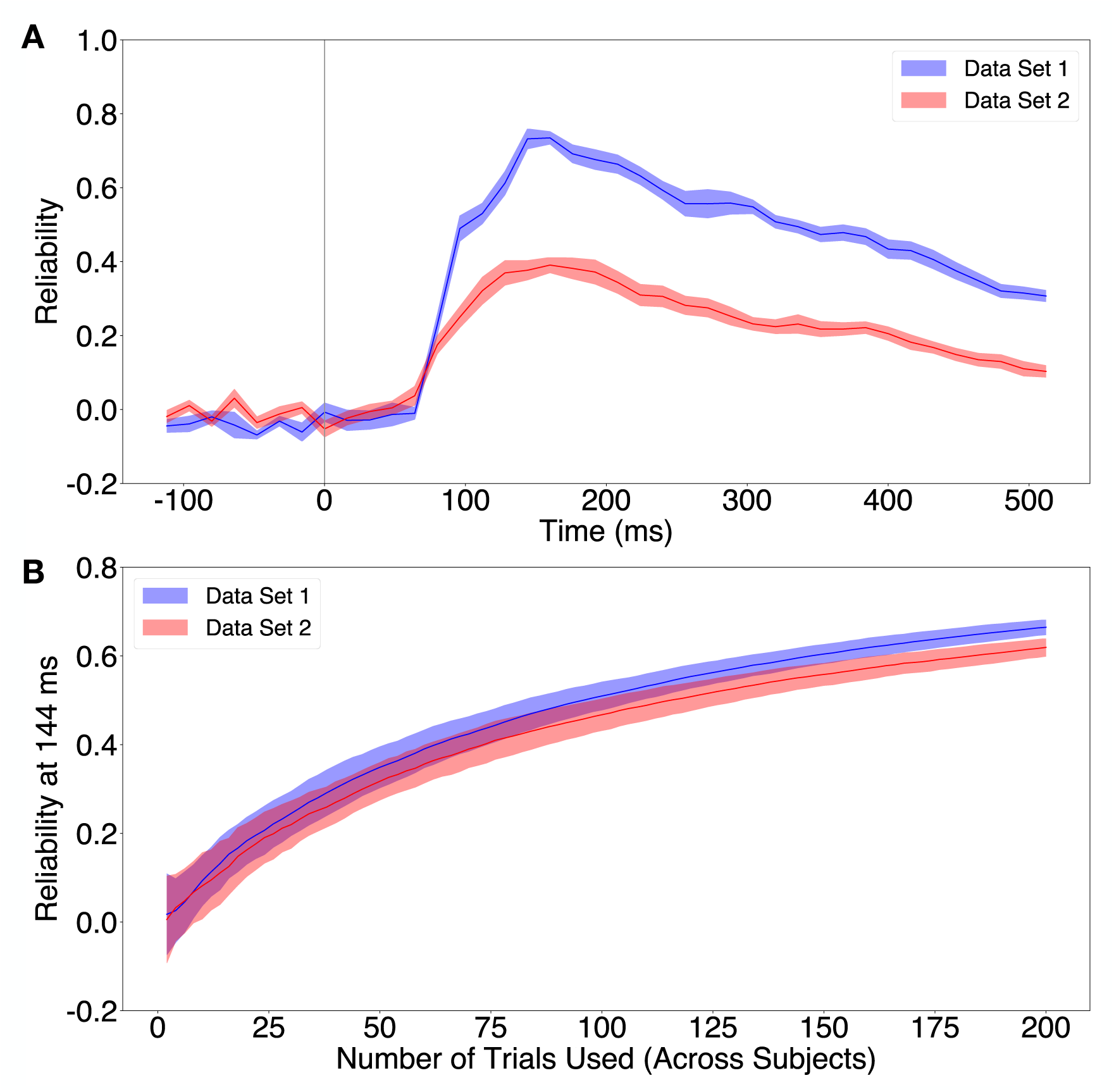
**A**: Test-retest reliability of EEG data averaged across electrodes as a function of time. The dark solid line indicates the average reliability across all subjects and the shaded region indicates the standard error of the mean across subjects. The vertical grey bar indicates time of stimulus onset. **B**: Test-retest reliability of EEG data as a function of number of trials in the data sets. The dark solid line indicates the average reliability across 100 random pseudo-data sets and the shaded region indicates the standard deviation across 100 random pseudo-data sets.

In order to fairly compare the reliability of both data sets, we controlled for the number of trials used in the reliability computation. Figure 2B shows how the reliability of the EEG time series at 144 ms varies as a function of the number of trials used in each data set. As expected, the reliability increases as more trials are used in each data set. Furthermore, the qualitatively similar curves computed from Data Set 1 and from Data Set 2 indicate that the reliability of both data sets are comparable.

### 3.2. Representational Dissimilarity Matrices

RDMs convey the dissimilarity of stimulus pairs based on features from neural data or from models. Figure 3 shows rank-normalized EEG RDMs whose stimuli are organized by category (*N* = 6) and, within category, by individual category exemplars (*N* = 12)—specifically, the average of the temporally resolved EEG RDMs over all time points. In these plotted RDMs, blue entries indicate low dissimilarity and red entries indicate high dissimilarity. Figures 3A, 3B and 3C are the time-averaged RDMs computed using Data Set 1, and Figures 3D, 3E and 3F are the time-averaged RDMs computed using Data Set 2. Comparing the RDMs computed for each data set and for each similarity metric, we observe similarities in terms of the high-level structure of each RDM. For example, all RDMs exhibited a strong block along the diagonal for the human faces (HF), indicating that the neural responses to these stimuli exhibited low within-category dissimilarity but were robustly differentiable from the neural signals associated with stimuli from other image categories. There was also another relatively strong block along the diagonal for animal faces (AF) and, to a lesser extent, the two inanimate categories (FV and IO).

**Figure 3:**
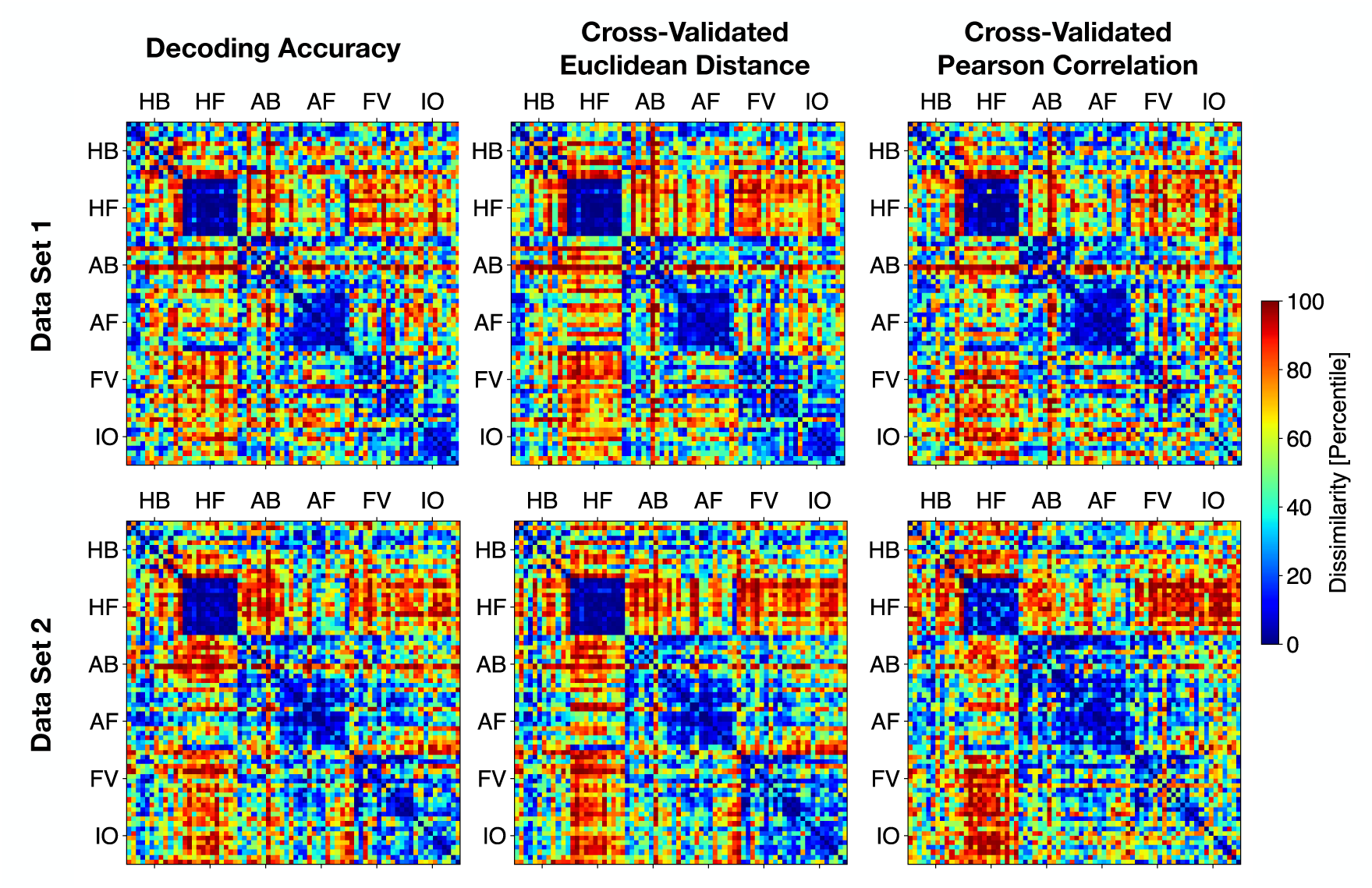
**Top**: rank-normalized EEG RDMs computed with Data Set 1. **Bottom**: rank-normalized EEG RDMs computed with Data Set 2. Blue indicates the lowest rank and red indicates the highest rank. **Left**: decoding accuracy metric. **Middle**: cross-validated Euclidean distance metric. **Right**: cross-validated Pearson correlation metric. These RDMs are the temporal average of RDMs computed at each time sample of the response with each similarity metric. The axes indicate the associated category of the stimulus. both human (HB and HF) categories, while blocking in the EEG RDMs more strongly implicated individual face (HF and AF) categories.

For comparison with the EEG RDMs, we also computed rank-normalized RDMs from representative layers of the CNN, as shown in Figure 4. As before, blue entries indicate low dissimilarity and red entries indicate high dissimilarity. We observe that the blocks along the diagonal corresponding to image categories become more distinct the deeper the layer is embedded in the neural network. For example, one can compare the RDMs from `pool2` and from `pool5` (Figure 4) and can observe that the blocks along the diagonal of the `pool5` RDM are qualitatively more apparent than those of the `pool2` RDM. At a high level, a major difference between the CNN RDMs and the EEG RDMs is that the CNN RDMs have large blocking across

**Figure 4:**
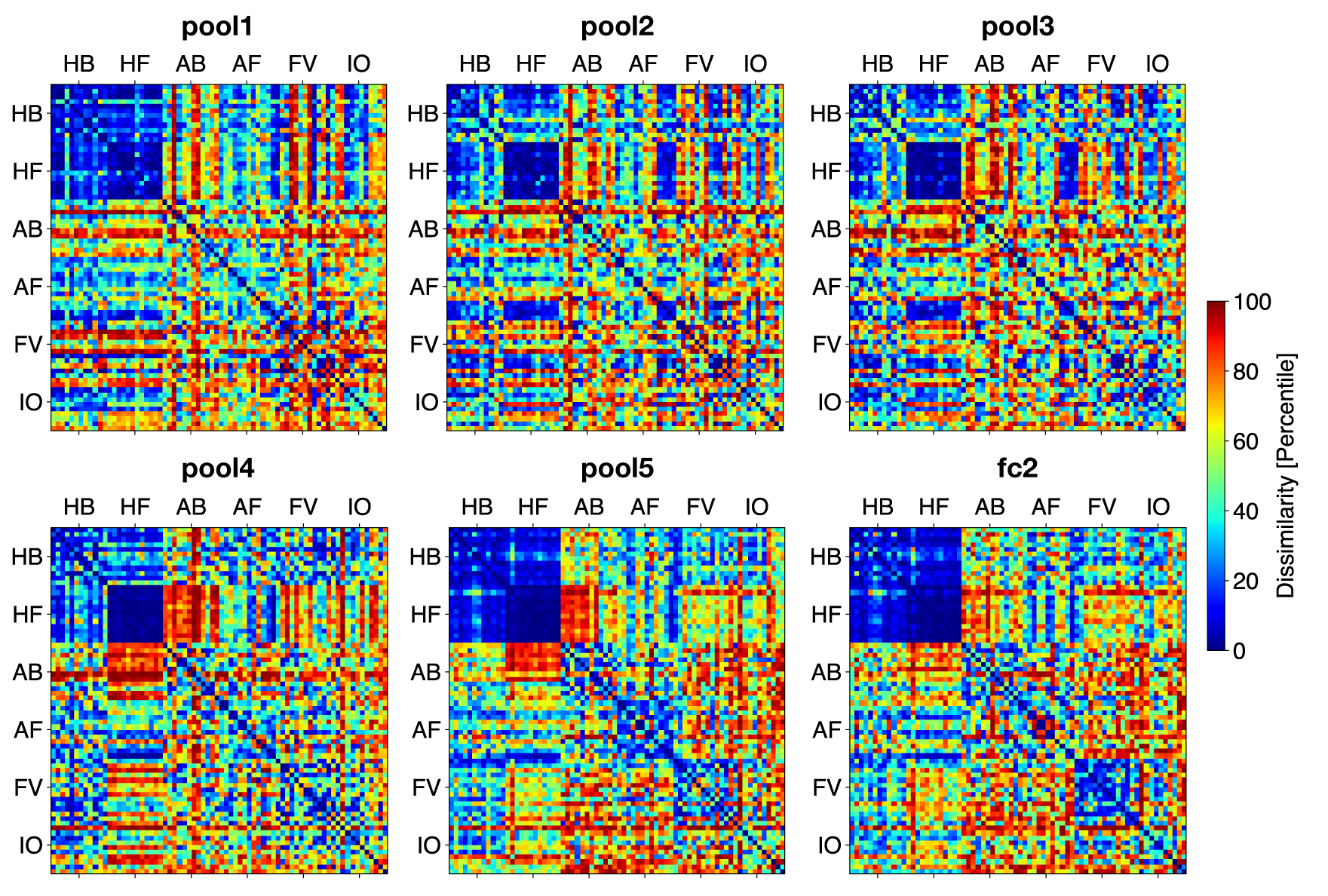
Rank-normalized RDMs computed using features in representative layers of the CNN. Blue indicates the lowest percentile ranked dissimilarity and red indicates the highest. The axes indicate the associated category of the stimulus.

### 3.3. Correlation Time Courses

#### 3.3.1. Comparing Per-Layer Correlation Time Courses

Taking advantage of the high temporal resolution of EEG data, we computed RDMs for each time sample of the response data and compared them to RDMs computed at various layers of the neural network. This allowed us to observe the ways in which representations from each layer of a neural network map on to representations at various time points in the neural response.

We analyzed the correlation between temporally resolved EEG RDMs and RDMs from the neural network model. We focus on EEG RDMs computed with the cross-validated Euclidean distance metric because this metric was shown to be a strong choice for RSA due to its reliability (Guggenmos et al., 2018). We also observed that EEG RDMs computed using this distance metric for both data sets were more reliable than those computed using other distance metrics (compare the noise ceiling in Figure 5 with those in Figures S1 and S2). Each layer of the neural network was associated with temporally resolved RDMs computed from each EEG data set. By superimposing the correlation time courses computed for each layer, we could observe the evolution of the relationship between the EEG RDMs and the neural network RDMs as a function of time and also as a function of neural network layer depth.

**Figure 5:**
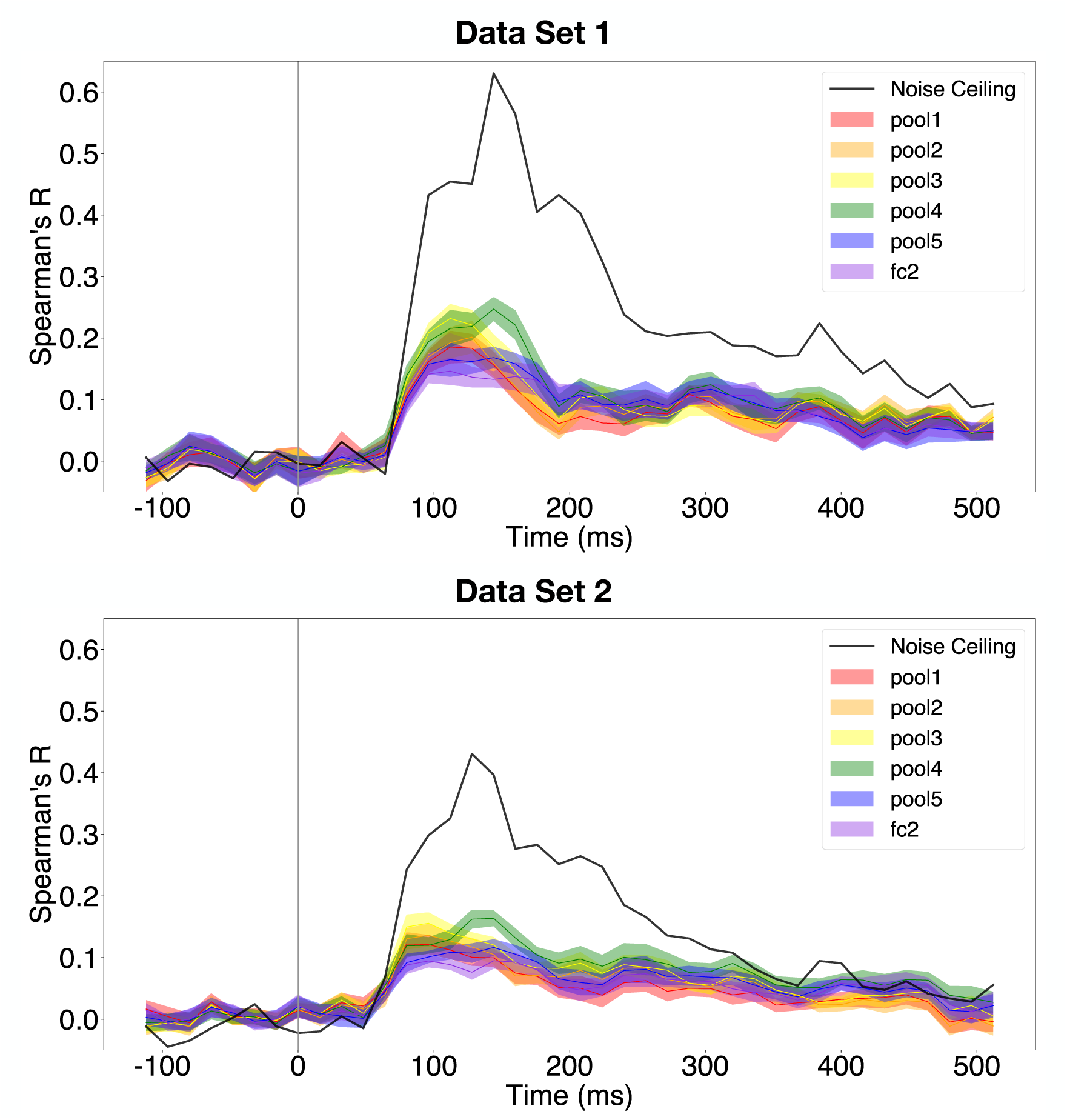
Correlation time courses for all layers using RDMs computed with the cross-validated Euclidean distance metric. Solid lines indicate the mean correlation across subjects and the shaded regions indicate the standard error of the mean across subjects. The vertical grey bar indicates time of stimulus onset. **Top**: Correlation time courses computed using Data Set 1. **Bottom**: Correlation time courses computed using Data Set 2.

Figure 5 shows how the correlations between RDMs computed from the model and from the two data sets varied with time. Each colour in Figure 5 represents a layer in the CNN that was used in the comparison with neural data. By observing the correlation time courses, we see that the time at which correlations started to deviate from zero was very similar for the representative layers in the neural network. However, the time at which the peak correlation occurred for each layer was later for layers that were deeper in the neural network. These qualitative observations can be observed in Figure 5, where each colour represents a layer in the neural network model. Importantly, these qualitative observations hold for both data sets. We quantitatively validate these observations in Sections 3.3.3 and 3.3.4 after showing how correlations vary when different similarity metrics are used.

Finally, we note that the peak correlation was approximately *R ≈* 0.25 for Data Set 1 and approximately *R ≈* 0.15 for Data Set 2. The corresponding correlation time course noise ceilings were much higher than the peak correlations reached by this neural network model, indicating that the low correlations were not the result of noisy neural data, but rather the result of a poor model fit.

#### 3.3.2. Comparing Metrics

We next compared correlation time courses across similarity metrics used to compute the EEG RDMs. In the next set of plots, each plot focusses on one neural network layer and overlays the correlation time courses for all three similarity metrics. This allows us to directly compare the three similarity metrics on the basis of how the correlation time courses vary with respect to one particular layer. Here, we focus on a representative layer in the neural network, namely `pool1`. This layer represents a shallow layer of the CNN. Figure 6A shows the correlation time course which compares RDMs computed from `pool1` with RDMs computed using all three methods. We observe that the correlation time courses were very similar if the decoding accuracy metric and the cross-validated Euclidean distance metric were used. Correlation time courses obtained using the Pearson correlation metric differed slightly from the other two time courses.

**Figure 6:**
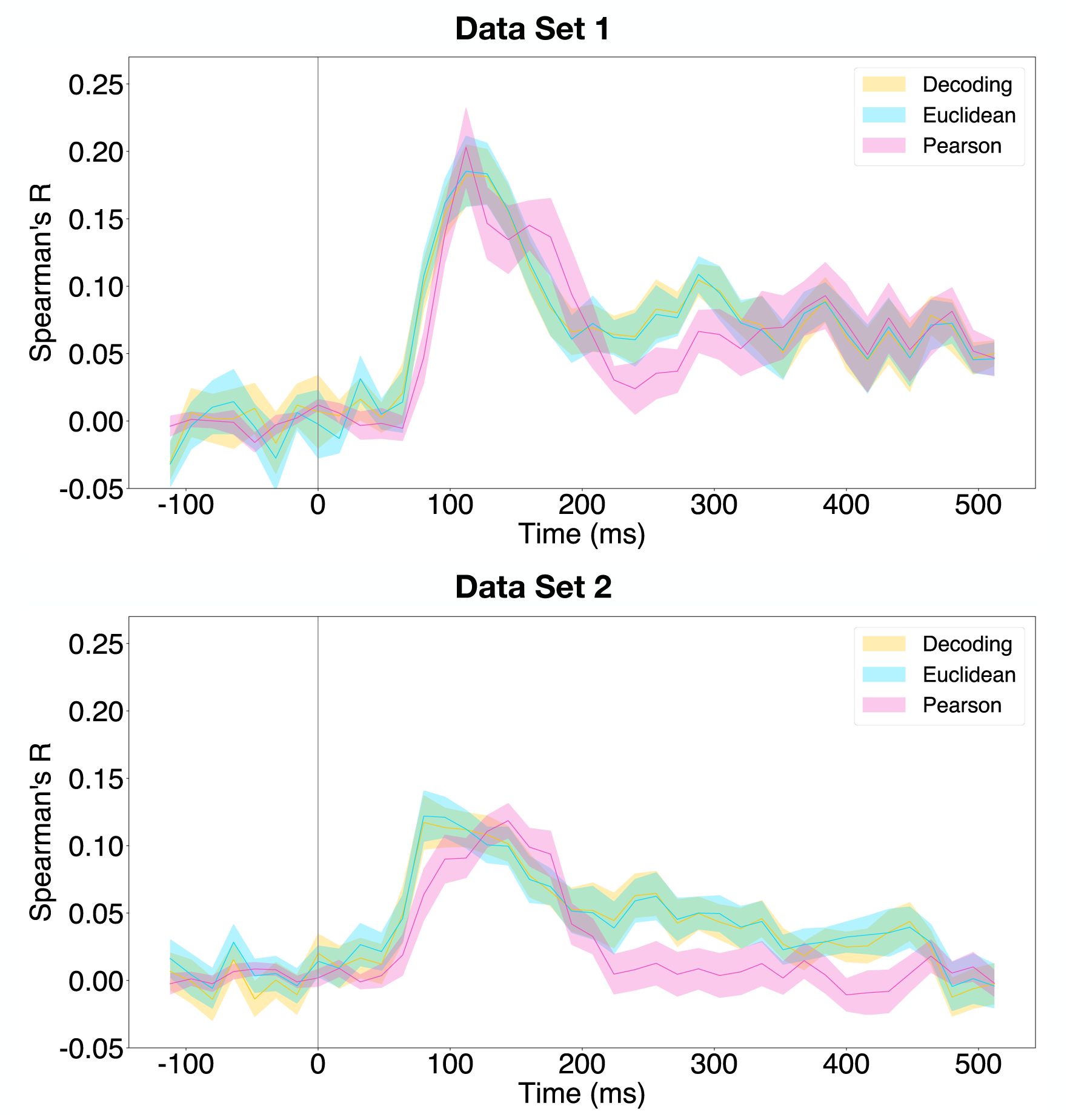
Correlation time course for `pool1` with respect to all three methods. Solid lines indicate the mean correlation across subjects and the shaded regions indicate the standard error of the mean across subjects. The vertical grey bar indicates time of stimulus onset. **Top**: Correlation time courses computed using Data Set 1. **Bottom**: Correlation time courses computed using Data Set 2.

The same analysis was performed on Data Set 2. Figure 6B is the correlation time course for `pool1`. A similar observation to that found in Data Set 1 can be found in these correlation time courses. In this data set, correlation time courses obtained by using RDMs computed with decoding accuracy and Euclidean distance as the similarity metric were similar. Similarly to the observation from the analyses of Data Set 1, we see that the time course obtained from the Pearson correlation differed slightly from the other two time courses.

#### 3.3.3. Quantification of Peak Correlation Timings for Each Layer and Similarity Metric

We assessed quantitatively how the time of peak model-EEG RDM correlation varied with increasing depth of the layers in the CNN. Hypothetically, if the layers of a neural network reflect the hierarchical organization of the brain’s visual areas, the time of peak correlation should increase as the depth of the neural network layer increases.

Depending on the similarity metric used to compute the RDMs from the neural data, we observed a positive relationship between the time of peak correlation and the depth of the layer in the CNN. Figure 7A shows how the time of peak correlation varies with CNN layer for Data Set 1. When EEG RDMs were computed using the decoding accuracy and Pearson correlation metric, the slope was significantly positive (*p <* 0.05), indicating that the time of peak correlation occurrence increased as a function of depth in the CNN. On the other hand, when the Euclidean distance was used, the slope was not significantly different from 0. Figure 7B shows how the time of peak correlation varies with CNN layer for Data Set 2. The analyses on this data set showed that there was a positive relationship between depth and time of peak correlation (*p <* 0.05) for EEG RDMs computed using the decoding accuracy and Euclidean distance metric. When the Pearson correlation metric was used, the slope was not significantly different from 0.

**Figure 7:**
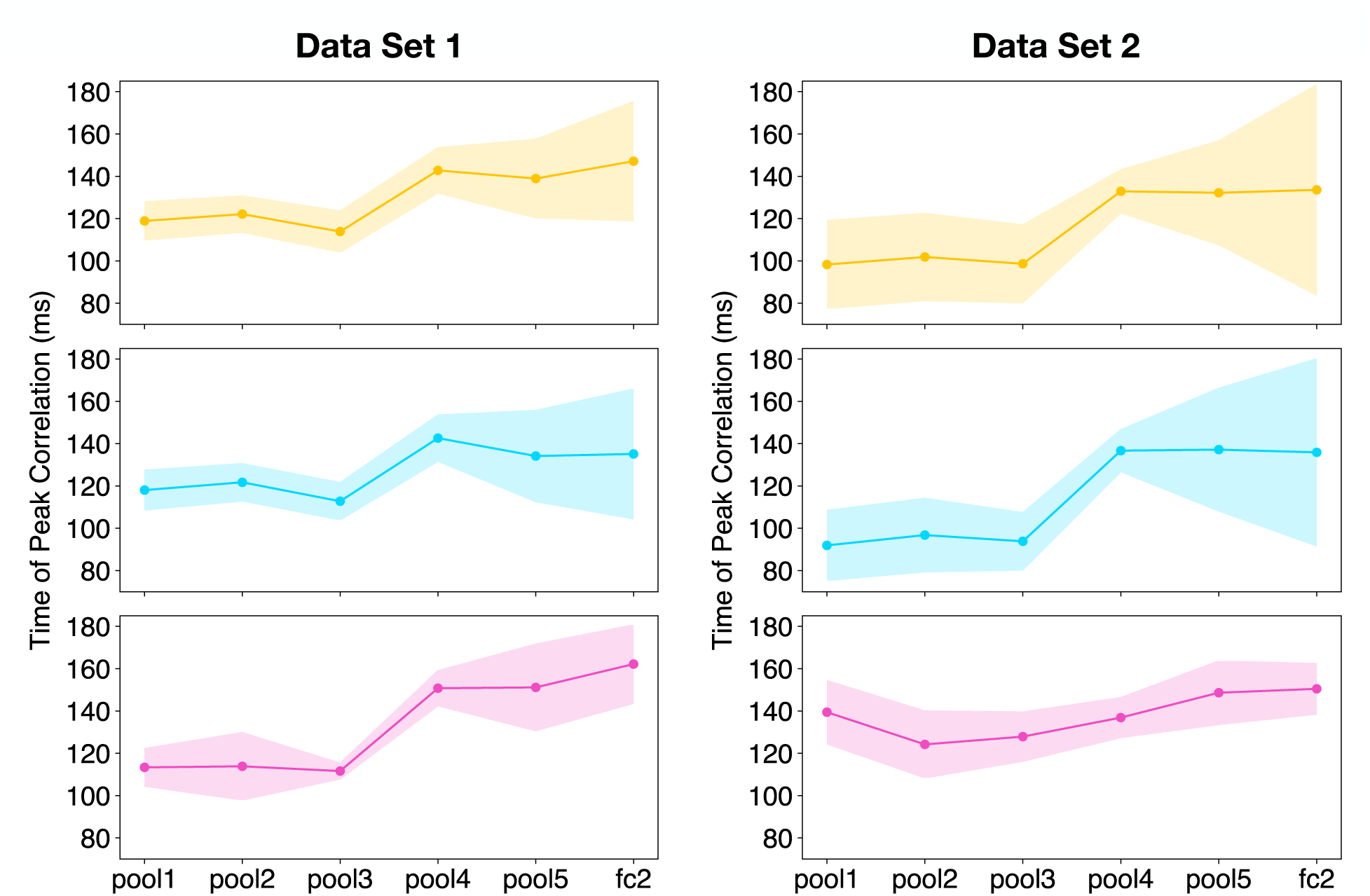
Time at which the peak correlation in the correlation time course occurs as a function of the layer in the neural network. Each colour represents different similarity metrics used to compute the RDMs. **Yellow**: decoding accuracy. **Blue**: Euclidean distance. **Pink**: Pearson correlation. Solid line indicates average time of peak correlation computed across 1000 bootstrap samples. Shaded regions indicate the standard error of the mean computed from 1000 bootstrap samples across subjects. **Left**: Using Data Set 1. Top: slope = 6.3, *p <* 0.05. Middle: slope not significantly different from 0. Bottom: slope = 11.3, *p <* 0.05. **Right**: Using Data Set 2. Top: slope = 8.62, *p <* 0.05. Middle: slope = 11.0, *p <* 0.05. Bottom: slope not significantly different from 0.

#### 3.3.4. Quantifying Time of Correlation Onset for Each Layer and Similarity Metric

Similar to the hypothesis that the time of peak correlation should vary with neural network layer depth, we expected that the correlation onset time should increase as the depth of the layer in the CNN increases. That is, when the correlation onset time is plotted against layers of increasing depth, the slope of the line of best fit should be positive. Concretely, we would observe a positive slope of the line that is fit between the correlation onset time and the depth of the neural network layer (DiCarlo et al., 2012).

We did not observe a positive relationship between the correlation onset time and the depth of the layer in the neural network model for either EEG data set. Figure 8A shows how the correlation onset time varies with the representative layers of the CNN for Data Set 1. In contrast to the peak correlation analyses, we did not observe a significant relationship between the time of correlation onset and the depth of the neural network layer for RDMs computed from any of the three similarity metrics (*p >* 0.05). Figure 8B similarly shows how the correlation onset time varies with the representative layers of the CNN for Data Set 2. The relationship we observed was the same as that from the analyses of Data Set 1.

**Figure 8:**
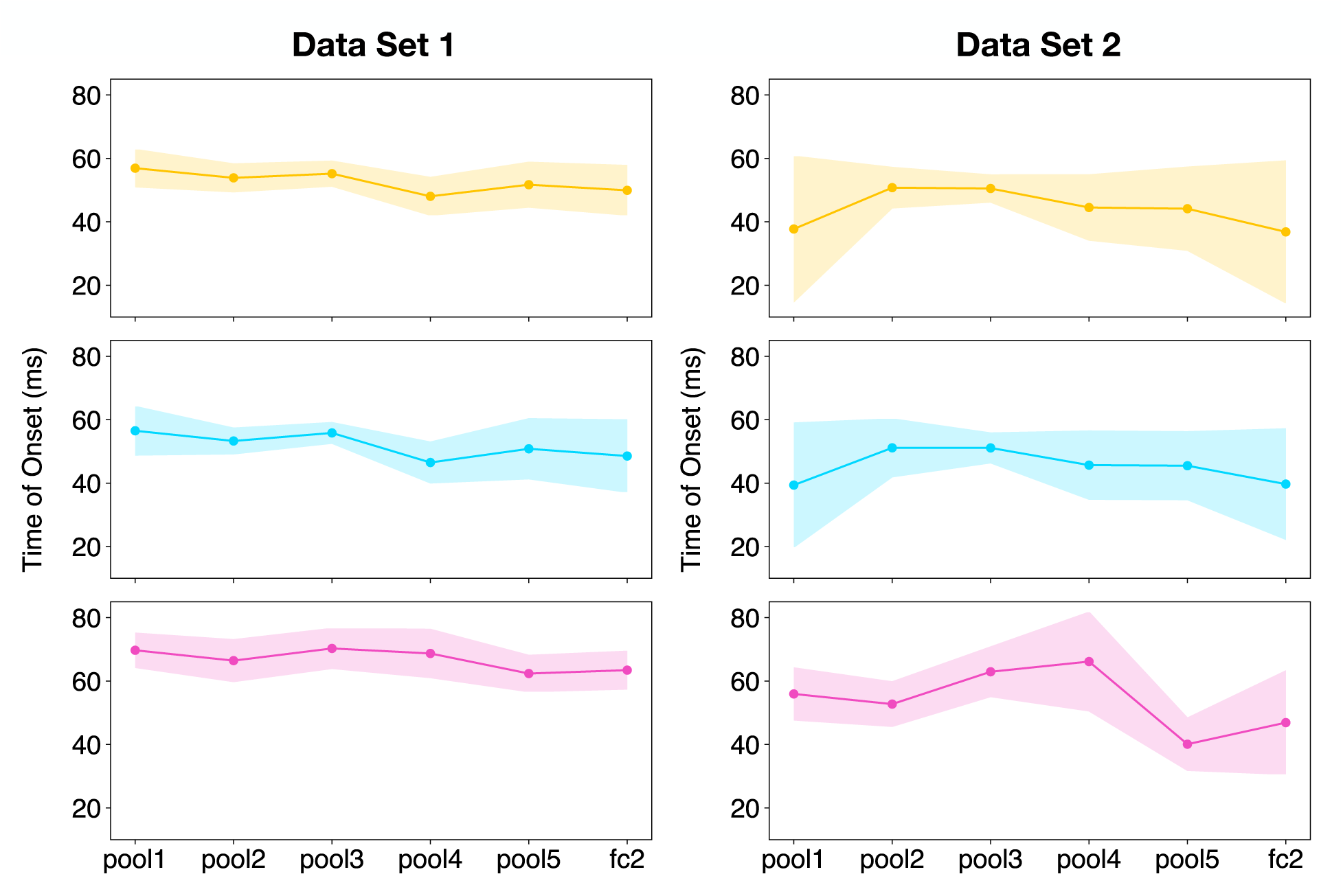
Time at which the correlation between RDMs computed from EEG data and RDMs computed from the neural network starts to increase. Each colour represents different similarity metrics used to compute the RDMs. **Yellow**: decoding accuracy; **Blue**: Euclidean distance; **Pink**: Pearson correlation. Solid lines indicate average time of peak correlation computed across 1000 bootstrap samples. Shaded regions indicate the standard error of the mean computed from 1000 bootstrap samples across subjects. **Left**: Using Data Set 1. Top, middle, bottom: slopes not significantly different from 0. **Right**: Using Data Set 2. Top, middle, bottom: slopes not significantly different from 0.

### 3.4. Correlation Time Courses from Optimal Linear Combinations of Features from the CNN

Linear reweighting of features from a model has previously been shown to improve correlations between a model RDM and human and monkey IT RDMs (Khaligh-Razavi and Kriegeskorte, 2014). Here we computed at each time point an optimal set of features which were linear combinations of the features from a particular layer of the CNN and then constructed RDMs using these synthesized features. Similar to the hypotheses described in the previous sections, we expected that deeper layers would provide stronger correspondences to EEG RDMs at later time points than those of shallow layers.

Here we focus on EEG RDMs that were computed using the cross-validated Euclidean distance metric for reasons described previously (see Section 3.3.1). Figure 9 shows how the correlations between the EEG RDMs computed with the cross-validated Euclidean distance metric and the RDMs computed from optimal feature combinations vary with time. Firstly, we observe that using optimally combined features from the CNN results in RDMs that are more correlated to EEG RDMs—the absolute Spearman’s R values are higher using the optimization procedure (compare Figure 5 with Figure 9).

**Figure 9:**
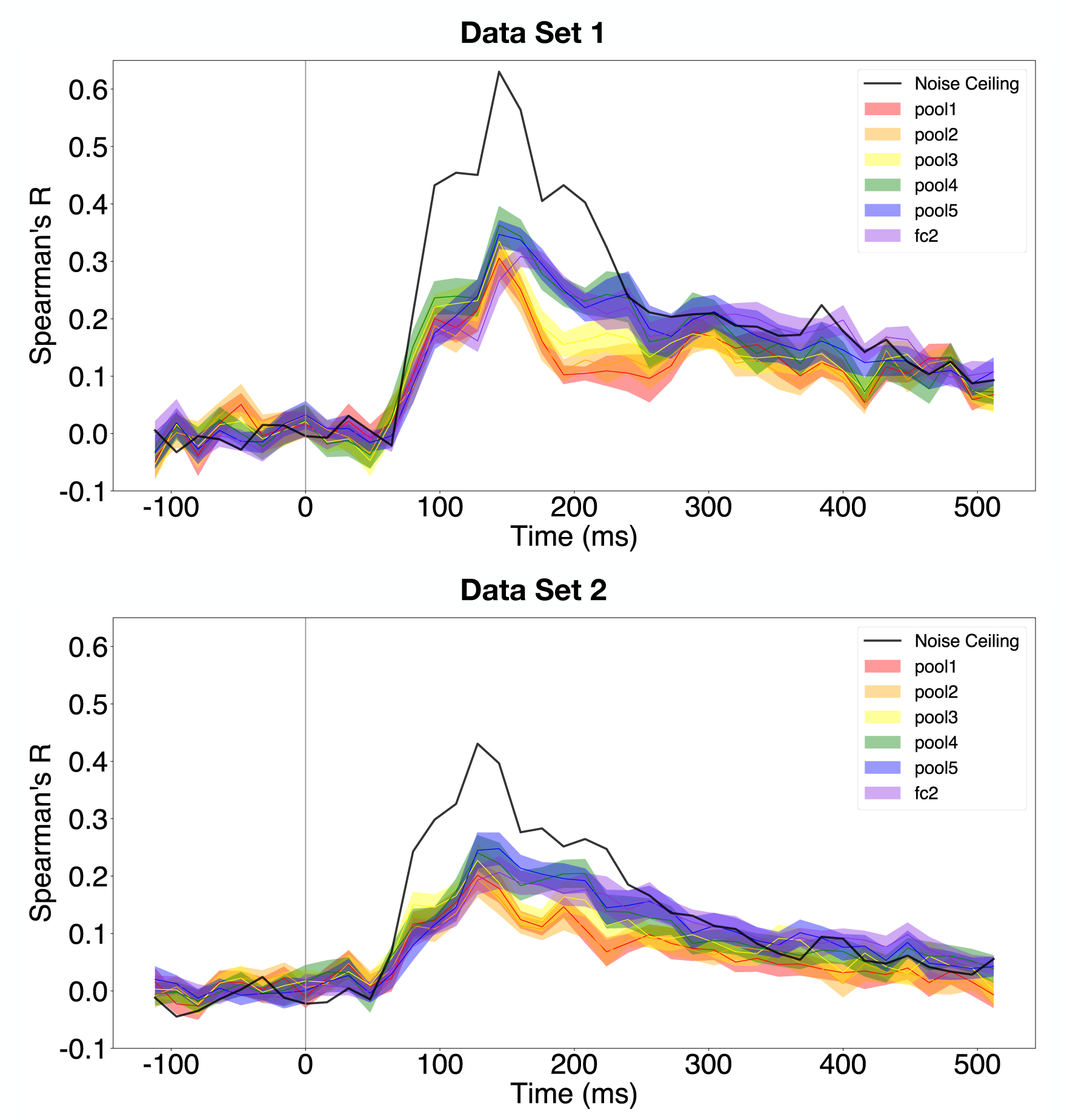
Correlation time courses for all layers using RDMs computed with the cross-validated Euclidean distance metric and the linear combination optimization procedure. Solid lines indicate the mean correlation across subjects and cross-validation folds and the shaded regions indicate the standard error of the mean across subjects. The vertical grey bar indicates time of stimulus onset. **Top**: Correlation time courses computed using Data Set 1. **Bottom**: Correlation time courses computed using Data Set 2.

Next we wanted to investigate the relationship between the layers of the CNN and the time at which the CNN layer RDMs are best able to correlate with EEG RDMs. We focus on response latencies between approximately 80 *−* 200 ms from stimulus onset because in macaque, inputs reach primary visual cortex at approximately 50 ms and reach IT cortex at approximately 100 ms (DiCarlo et al., 2012). Thus, applying the 5*/*3 conversion rule from macaque to humans (Schroeder et al., 2004), these time values convert to approximately 83 ms and 167 ms. Figure 10 shows how the noise-ceiling normalized correlations vary between 80 ms and 200 ms. Qualitatively, we see that deeper layers seem to provide a better correspondence to EEG RDMs at time points later than approximately 176 ms.

**Figure 10:**
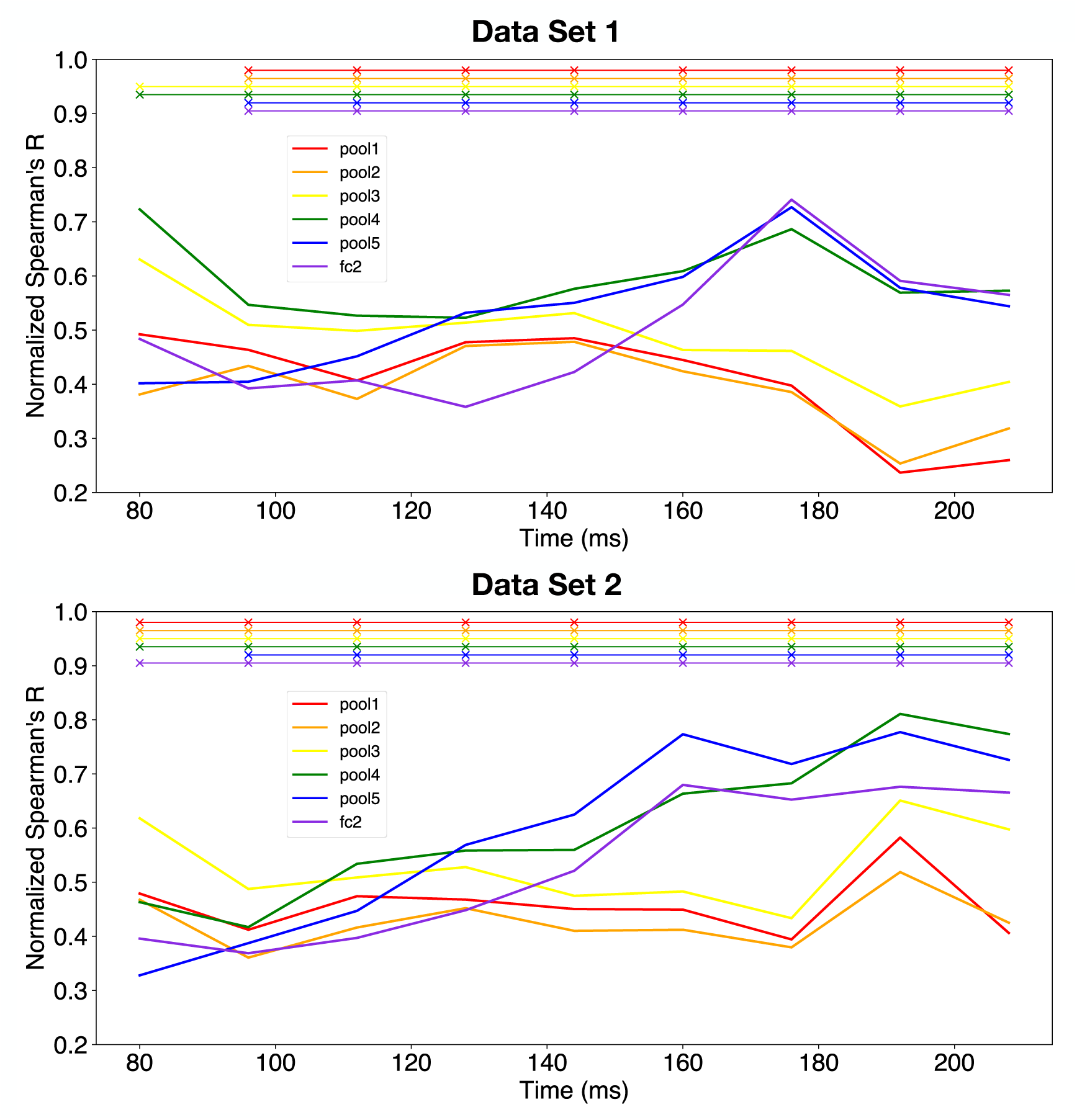
Noise-ceiling normalized correlation time courses from 80 *−* 200 ms post-stimulus onset for all layers using RDMs computed with the cross-validated Euclidean distance metric and the linear combination optimization procedure, which were computed by dividing the absolute Spearman’s R by the noise ceiling value at each time point. Solid lines indicate the mean correlation across subjects and cross-validation folds and the *×* symbols at the top of the figure indicate statistically significant correlations (*p <* 0.05, Bonferroni corrected for six comparisons). **Top**: Noise-ceiling normalized correlation time courses computed using Data Set 1. **Bottom**: Noise-ceiling normalized correlation time courses computed using Data Set 2.

To show that this is the case quantitatively, we plot and analyze the difference in noise-ceiling normalized correlations between `fc2` and every other layer between 80 ms and 200 ms. Figure 11 shows the time-varying differences in correlation between the noise-ceiling normalized correlation time courses from all the layers with respect to `fc2`. We see that the difference between `pool1` and `fc2` are significantly positive in the later time points indicating that the noise-ceiling normalized correlation is larger for `fc2` in those time points (*p <* 0.05, Bonferroni corrected for multiple comparisons). A similar observation can be found for `pool2` vs. `fc2`. This effect can also be observed when the decoding accuracy metric is used to compute EEG RDMs (see Figure S10), but is not significantly observed when the cross-validated Pearson correlation metric is used (see Figure S13).

**Figure 11:**
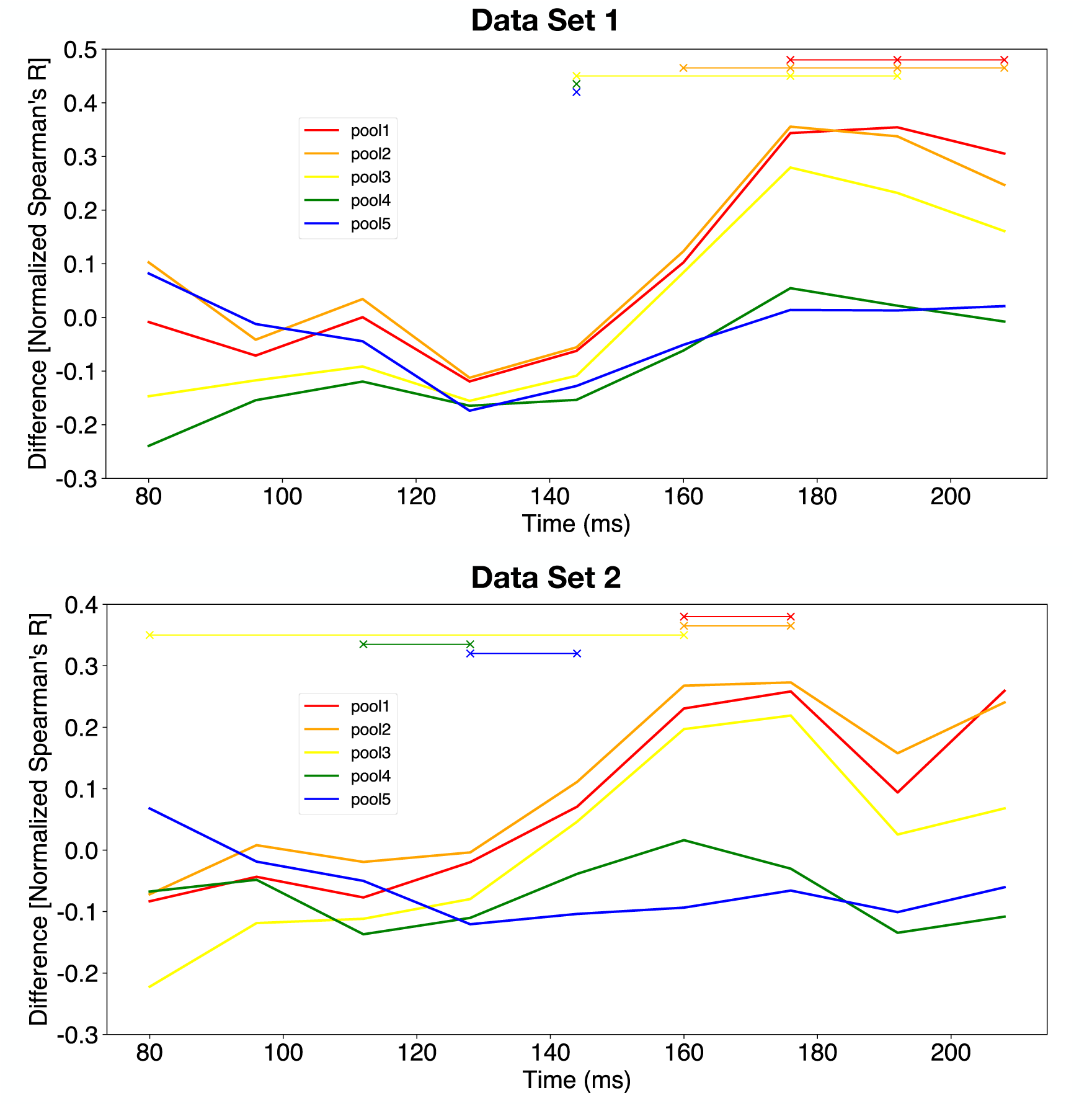
Differences in noise-ceiling normalized correlation time courses from 80 *−* 200 ms after stimulus onset for all layers with respect to `fc2` using RDMs computed with the cross-validated Euclidean distance metric and the linear combination optimization procedure. These were computed by subtracting the noise-ceiling normalized correlations of every other CNN layer from that of `fc2`. Positive values indicate that the correlation value for `fc2` is larger than that of the respective layer. Similarly, negative values indicate that the correlation value for `fc2` is smaller than that of the respective layer. Solid lines indicate the mean correlation across subjects and cross-validation folds and the *×* symbols at the top of the figure indicate statistically significant differences (*p <* 0.05, Bonferroni corrected for five comparisons). **Top**: Differences in noise-ceiling normalized correlation time courses computed using Data Set 1. **Bottom**: Differences in noise-ceiling normalized correlation time courses computed using Data Set 2.

### 3.5. Correlation Time Courses from Optimal Element-Wise Scaling of Features from the CNN

Due to the computationally intense nature of the linear combination optimization procedure, an element-wise scaling optimization procedure was implemented. At each time point, we computed an optimal scalar value for each feature in a particular CNN layer. As before, we expected that deeper layers would provide a stronger correspondence to later time points in the EEG responses and that shallow layers would provide a stronger correspondence to earlier time points.

Here we also focus on the EEG RDMs computed using the cross-validated Euclidean distance metric. Figure 12 shows the correlation time course obtained by finding a set of weights that scaled each CNN feature at each time point. Qualitatively, we also observed that with the optimization, the absolute correlation values are higher than those computed without the optimization procedure (compare Figure 12 with Figure 5). However, as expected, the correlation values obtained using the element-wise optimization procedure were lower than those computed using the linear combination optimization procedure (compare Figure 12 with Figure 9).

**Figure 12:**
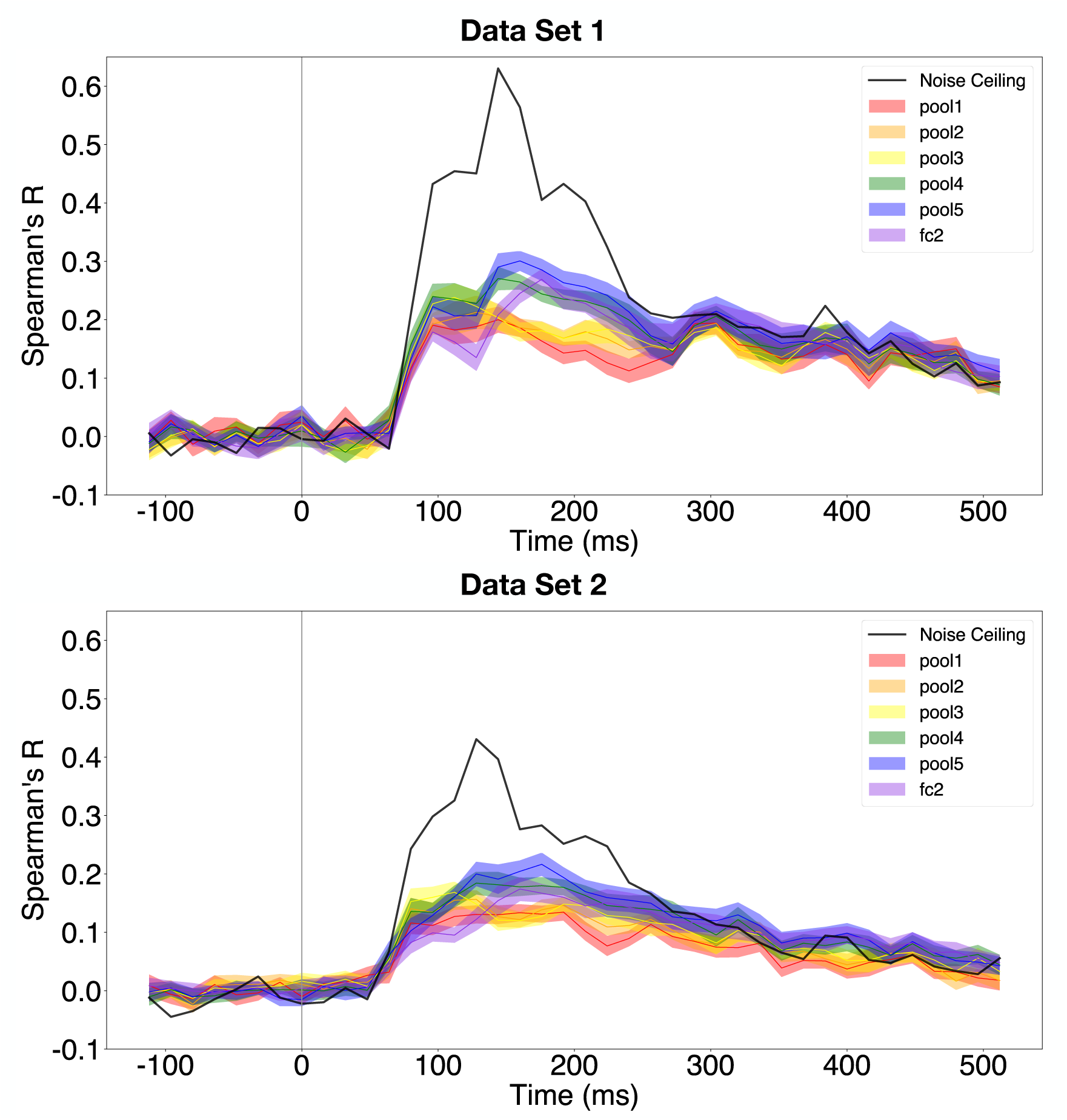
Correlation time courses for all layers using RDMs computed with the cross-validated Euclidean distance metric and the element-wise scaling optimization procedure. Solid lines indicate the mean correlation across subjects and cross-validation folds and the shaded regions indicate the standard error of the mean across subjects. The vertical grey bar indicates time of stimulus onset. **Top**: Correlation time courses computed using Data Set 1. **Bottom**: Correlation time courses computed using Data Set 2.

We also studied the relationship between the layers of the CNN and the time at which the layer RDM maximally corresponded to the EEG RDMs in the time range of approximately 80 ms and 200 ms. Qualitatively, we observed from Figure 13 that deeper layers tended to have a higher noise-ceiling normalized correlation with EEG RDMs at later time points (i.e. approximately 176 ms). We quantified this observation by analyzing the differences in the noise-ceiling normalized correlation between the representative layers of the CNN and `fc2` as seen in Figure 14. Similar to the observation described in Section 3.4, we observed that the difference between the noise-ceiling normalized correlations of `pool1` and `fc2` is significantly positive, indicating that the deeper layer provides a stronger correspondence than the shallow layer to later time points in the EEG responses (*p <* 0.05, Bonferroni corrected for multiple comparisons). As before, this effect is present when the decoding accuracy metric is used (see Figure S16), but cannot be observed when the cross-validated Pearson correlation metric is used (see Figure S19).

**Figure 13:**
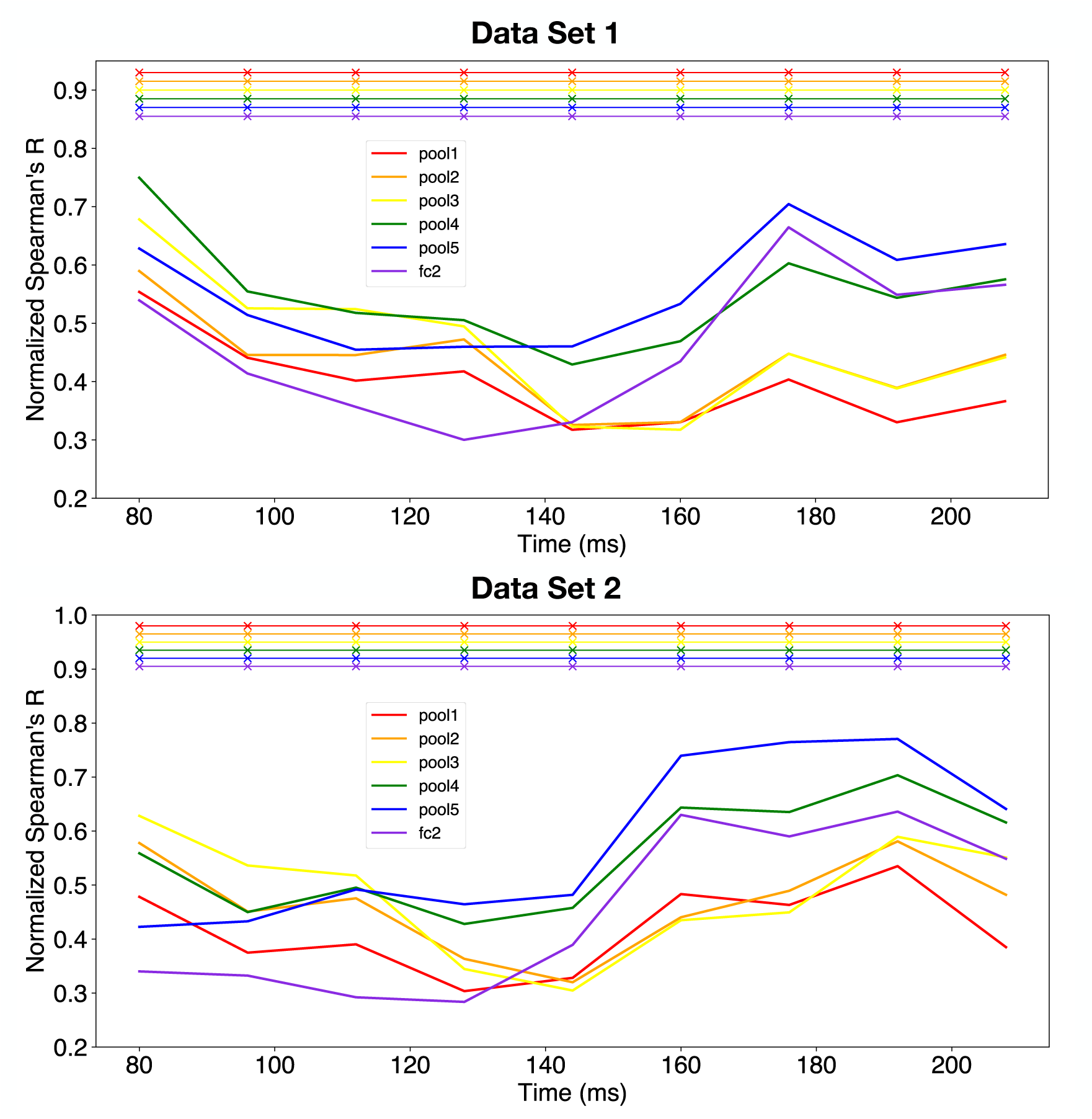
Noise-ceiling normalized correlation time courses for all layers using RDMs computed with the cross-validated Euclidean distance metric and the element-wise scaling optimization procedure, which were computed by dividing the absolute Spearman’s R by the noise ceiling value at each time point. Solid lines indicate the mean correlation across subjects and cross-validation folds and the *×* symbols at the top of the figure indicate statistically significant correlations (*p <* 0.05, Bonferroni corrected for six comparisons). **Top**: Noise-ceiling normalized correlation time courses computed using Data Set 1. **Bottom**: Noise-ceiling normalized correlation time courses computed using Data Set 2.

**Figure 14:**
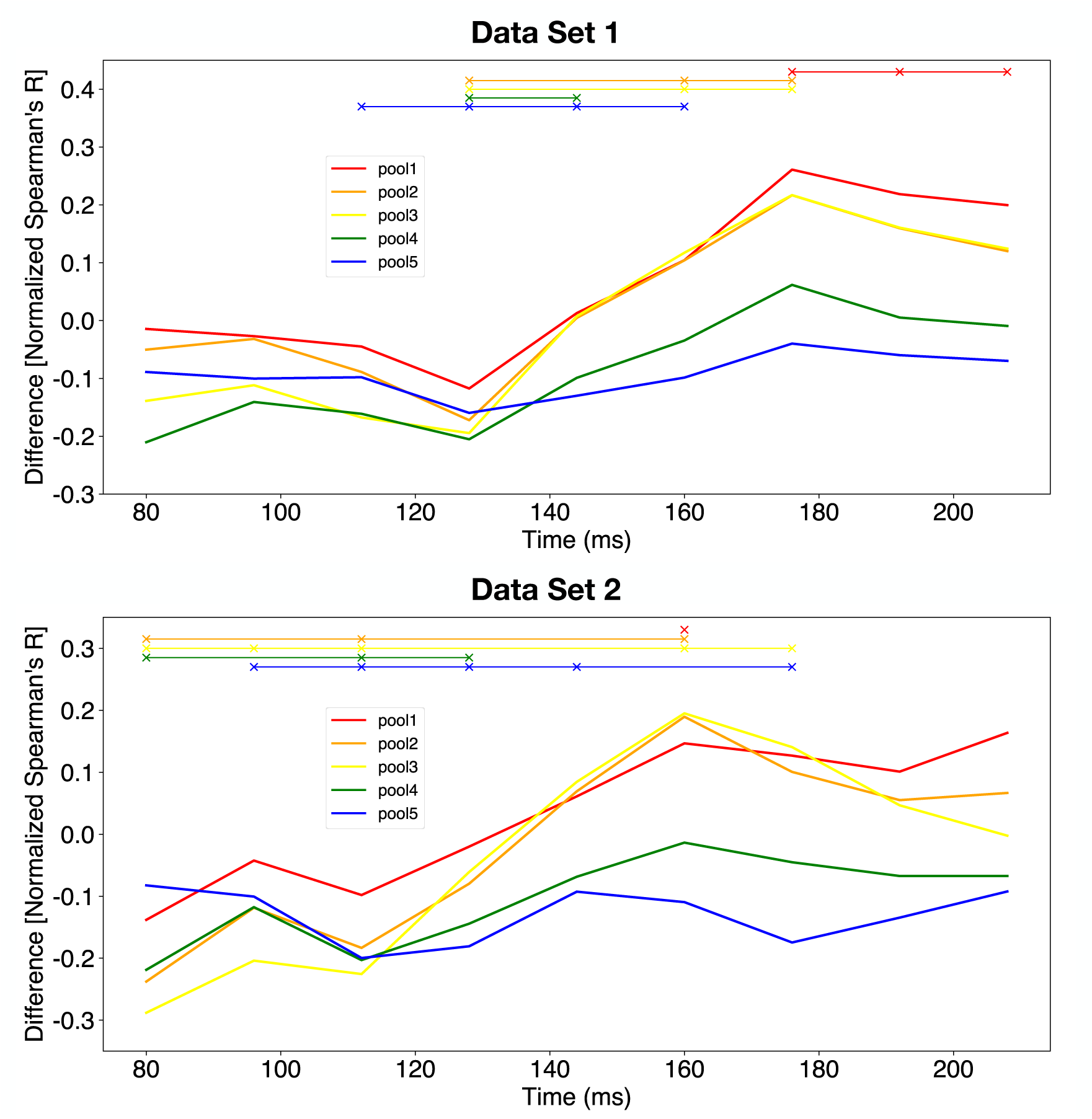
Differences in noise-ceiling normalized correlation time courses for all layers with respect to `fc2` using RDMs computed with the cross-validated Euclidean distance metric and the element-wise scaling optimization procedure. These were computed by subtracting the noise-ceiling normalized correlations of every other CNN layer from that of `fc2`. Positive values indicate that the correlation value for `fc2` is larger than that of the respective layer. Similarly, negative values indicate that the correlation value for `fc2` is smaller than that of the respective layer. Solid lines indicate the mean correlation across subjects and cross-validation folds and the *×* symbols at the top of the figure indicate statistically significant differences (*p <* 0.05, Bonferroni corrected for five comparisons). **Top**: Differences in noise-ceiling normalized correlation time courses computed using Data Set 1. **Bottom**: Differences in noise-ceiling normalized correlation time courses computed using Data Set 2.

## 4. Discussion

### 4.1. Summary

Using RDMs computed from representative layers of a CNN, we assessed the extent to which they correlated with RDMs computed from EEG data using various similarity metrics. In addition, we assessed the reproducibility of the results across EEG data sets. From a high-level perspective, we tested the hypothesis that RDMs computed using early layers in the CNN would strongly correlate with EEG RDMs computed at early time samples of the response. This hypothesis has previously been proposed and is based on the idea that features extracted in early layers of a CNN would be very similar to features extracted in early visual cortex (Güçlü and van Gerven, 2015; Cichy et al., 2016). As more information is processed over time, more complex features would be extracted in higher visual cortex, which would presumably correspond well with the more complex features extracted in the higher layers of the CNN.

We investigated how the time of peak correlation varied as a function of depth in the neural network model. In particular, we observed that when the decoding accuracy metric was used to construct EEG RDMs, there was a statistically significant positive relationship between the depth of the layer and the time at which the peak correlation occurs for both EEG data sets. This finding suggests a hierarchical correspondence between the neural network model and the temporal dynamics in the EEG data, as previously reported (Güçlü and van Gerven, 2015; Cichy and Pantazis, 2017). However, the positive relationship was found for the Pearson correlation metric only for Data Set 1 and for the Euclidean distance metric only for Data Set 2. Therefore, the decoding accuracy metric may be more robust.

We also studied how the model-EEG RDM correlation onset time varied as a function of depth in the neural network model with an expectation that onset times, like peak correlation times, should be delayed for deeper network layers. However, in both data sets and across all three similarity metrics, we found no significant relationship between the correlation onset time and the depth of the neural network layer.

Finally, we provided two different optimization procedures which resulted in stronger correlations between the layer RDMs and the EEG RDMs. The linear combination optimization procedure learned features that were optimal linear combinations of the CNN features and the element-wise scaling optimization procedure learned weights that scaled the existing CNN features. The former procedure had many more weights to optimize for and therefore required much longer computation time than that of the latter procedure. By analyzing the noise-ceiling normalized correlation values between 80 ms and 200 ms, we observed that depending on the similarity metric used to compute EEG RDMs, RDMs computed from deeper layers provided a significantly stronger correspondence to EEG RDMs computed at later time points. However, we did not observe that shallow layers corresponded better than deeper layers to earlier time points—shallow layers such as `pool1` provided correspondences to early time points that were *similar to* (i.e. not significantly different from) those of deeper layers such as `fc2` at early time points.

### 4.2. Hierarchical Correspondence in Time Between CNN Representations and EEG Responses

We observed that a correspondence between time of peak correlation and depth of CNN layer depended on both the similarity metric and the specific data set used. Moreover, this observation was supported by our findings when both the linear combination and element-wise scaling optimization procedures were used, where deep layers corresponded better than shallow layers to later time points in the EEG responses for specific similarity metrics. Thus, these results partially corroborate previous neuroimaging findings of a spatio-temporal correspondence between feed-forward CNNs and neural responses (Seeliger et al., 2017; Cichy et al., 2016).

Cichy et al. (2016) reported that the time of peak correlation increased with the depth of the layer in the CNN, using RSA and decoding accuracy as the similarity metric. This is consistent with the observations that we report here for the decoding accuracy similarity metric across both data sets. It is important to note that their peak correlation values are relatively low—the maximum correlation value of the correlation time courses reported in Cichy et al. (2016) are approximately 0.15. The peak correlation value that we report here is approximately 0.25 for Data Set 1 and approximately 0.2 for Data Set 2, without using the optimization procedures. These values, however, are well below the noise ceiling, which was not reported in Cichy et al. (2016).

Correlation values achieved without reweighting of CNN features were relatively low. However, when the optimization procedures were used, these correlation values were improved upon substantially. We also note that both the correlations achieved using the optimization procedures and the noise ceiling are high compared to those of neural data acquired using other imaging modalities such as fMRI (Khaligh-Razavi and Kriegeskorte, 2014). This could suggest that by using EEG responses, models could more easily be compared among each other.

Seeliger et al. (2017) used a different approach to address the correspondence between CNN layers and MEG responses. Instead of using RSA, the authors measured source-localized activity and performed a regression analysis using features of a different CNN model and current densities in different brain areas. They demonstrated hierarchical correspondence by showing that features from shallow layers of the CNN start to have stronger correlations (at least 0.3) with neural responses early in time, and that features of deeper layers of the CNN start to have stronger correlations with neural responses later in time.

### 4.3. Correspondence of Deep Layers of the CNN to EEG Responses at Early Time Points

In Section 3.3.4, we observed that the correlation onset for all representative layers was simultaneous. A similar finding regarding correlation onset times for RDMs was also reported in the supplemental information of Cichy et al. (2016), where the onset times were very similar across all the layers in their CNN (cf. Suppl. Table 2 of Cichy et al. (2016)), but was not commented on. By using a regression method similar to that used in Seeliger et al. (2017) instead of RSA, Ramakrishnan et al. (2017) reported that variance explained by summary statistics from various layers of a neural network starts to increase from zero at similar response latencies. Seeliger et al. (2017) made quantitative estimates of onset time based on the time at which a correlation of at least 0.3 occurred. We measured the time at which the correlation first started to rise. Given that we observed no hierarchical correspondence in our onset time measurements, but did in our peak correlation time measurements, it is thus likely that their onset metric is more related to our peak measurement. Each of these observations is inconsistent with expected increasing delays in a strictly hierarchical feed-forward model, where previous neural measurements show approximately 10-ms delays between visual areas (DiCarlo et al., 2012). One expects that these delays accumulate as one progresses from early to higher cortical areas. Therefore, although we observed a hierarchical correspondence for the time of peak correlation between layers of a CNN and EEG responses, the fact that our observed onset times were constant across CNN layers suggests only partial support for the hypothesis of a hierarchical correspondence.

A similar observation resulted from using the optimization procedures. Although we observed that depending on the similarity metric used to compute EEG RDMs, deeper layers corresponded better to later time points than did shallow layers, we did not observe the opposite—shallow layers did *not* correspond better than deeper layers to early time points.

A possible explanation for these phenomena is related to the uncontrolled nature of the stimulus set. Figure 12 of Kaneshiro et al. (2015b) shows that even after averaging images within an image category, it is possible to discriminate between categories due to the different categories being pixel-wise consistent within category, but different between category. This means that low-level features are highly correlated with high-level category. In fact, it was shown previously that when RDMs were computed using low-level information such as silhouettes, luminance and the V1 model, the human face category is highly distinct from other stimulus categories. This was evidenced by the strong human face block along the diagonal of the RDM (cf. Figure 6 of Kriegeskorte et al. (2008a)). This stimulus set structure suggests that features computed early in time (such as low-level features) can already provide information about image category. Conversely, deeper layers in the neural network may also be sensitive to low-level features that are correlated with category, which could drive the early onset of correlations. These phenomena also provide some explanation for the qualitatively similar RDMs computed from `pool1` and from deeper layers including `pool5` and `fc2` (see Figure 4). This suggests that in order to be able to differentiate shallow and deep CNN layer correspondences at early time points, stimuli that are used in the EEG experiments should be designed such that image category information should not be decodable using low-level image features.

### 4.4. Cross-Laboratory Consistency

Through reliability analysis, we observed that the reliability of both EEG data sets was comparable. Importantly, the fundamental observations (shift in time of peak correlation and constant correlation onset time) were consistent across data sets, especially for the decoding accuracy metric. The observations from the two optimization procedures were only somewhat consistent across both data sets, most likely due to the fact that EEG RDMs computed from Data Set 2 were less reliable than those computed from Data Set 1 (see, for example, the noise ceiling in Figure 5). This provides some reassurance that observations reported using this analysis framework and data modality can be reproduced in two independent laboratories.

### 4.5. Limitations and Future Directions

Future work needs to address two issues that were raised here. Firstly, we observed that the noise ceilings shown in Figures 5, 9 and 12 were high and higher than the achieved correlations even after the optimization procedure was performed. Past research using single-unit physiology has found a relatively small gap between the achieved variance explained and the noise ceiling (Cadena et al., 2019; Yamins et al., 2014). It is possible that the gap could be narrowed by using a different model class. A fundamental difference between the ventral visual stream and the CNN model used in this work is the presence of multiple feedback pathways between visual cortical areas (Gilbert and Li, 2013). Developing models whose architectures incorporate these connectivity patterns may prove to provide a better fit to the neural data. For example, a recurrent convolutional neural network has been shown to successfully model time-resolved single-unit recordings in macaque V4 and IT (Nayebi et al., 2018).

RSA is an intuitive method to visualize the similarity of two stimulus representations. However, our observations indicated that depending on the similarity metric used to compute EEG RDMs, different interpretations of the data may result. This was evidenced by the fact that if the Euclidean distance metric was used on Data Set 1 and if the Pearson correlation metric was used on Data Set 2, a non-significant positive relationship between time of peak correlation and depth of layer in the CNN resulted. The positive relationship was found for both data sets only if the decoding accuracy metric was used. A similar observation was found when both optimization procedures were performed. Deeper layer RDMs provided better correspondence than shallow layer RDMs when the EEG RDMs were computed using the cross-validated Euclidean distance. In particular, we noticed that when the cross-validated Pearson correlation metric was used to compute EEG RDMs, there was a non-significant correspondence between deeper layer RDMs and time. In the future, the observations presented here could be corroborated or rejected using other model comparison techniques. For example, one could develop linear models mapping the model responses to neural responses, which quantitatively improves the quality of the comparisons. The only issue that may arise is overfitting to a small stimulus set, so it is imperative to have a larger stimulus set (i.e. on the order of 10^3^) than that used in the current work when using this model comparison technique (Cadena et al., 2019). Finally, more careful design of stimuli would be beneficial in order to manipulate the correlations between low-level features and category-level information.

## Supplementary Material

**Figure S1:**
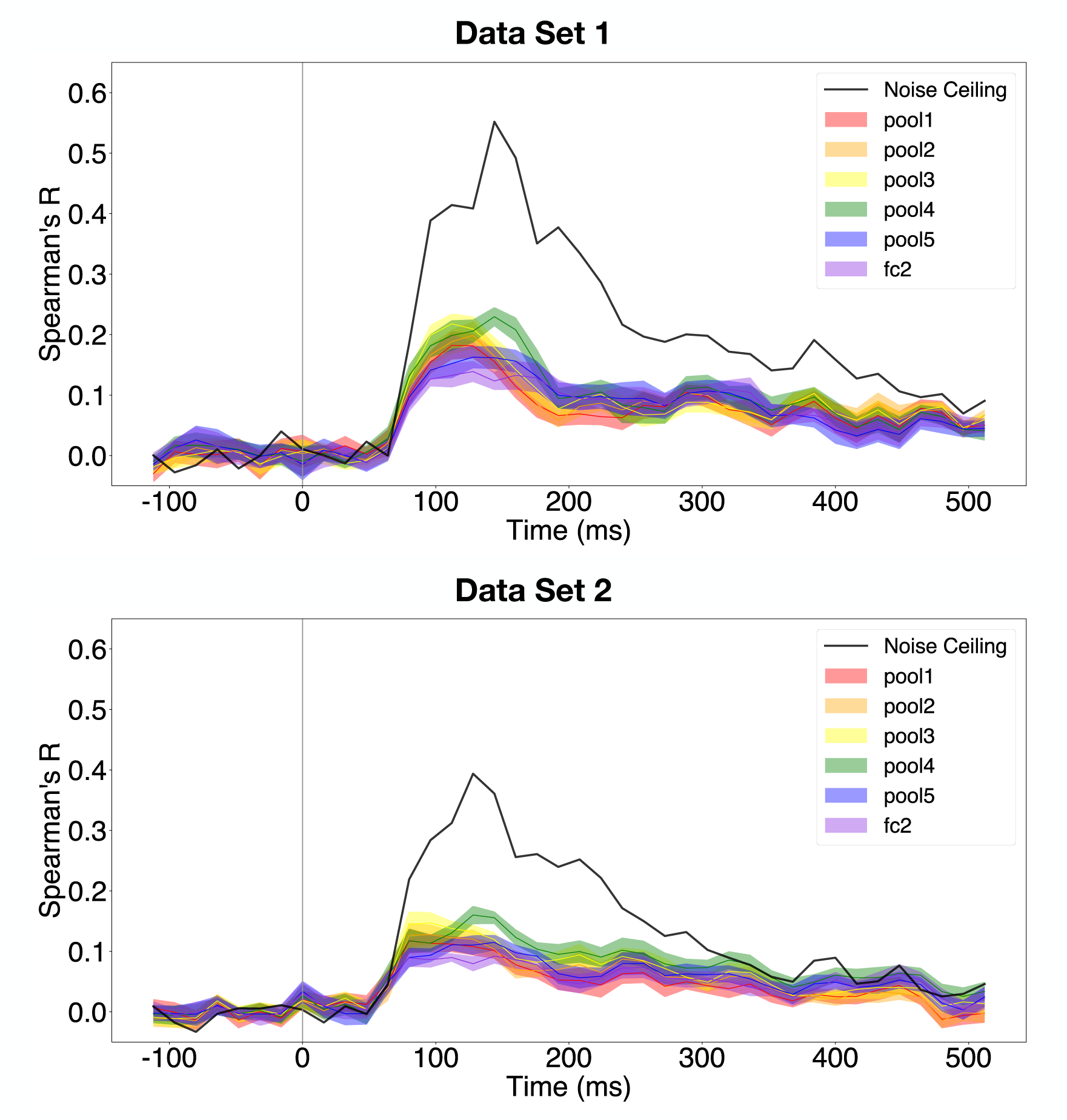
Correlation time courses for all layers using RDMs computed with the decoding accuracy metric. Solid lines indicate the mean correlation across subjects and the shaded regions indicate the standard error of the mean across subjects. The vertical grey bar indicates time of stimulus onset. **Top**: Correlation time courses computed using Data Set 1. **Bottom**: Correlation time courses computed using Data Set 2.

**Figure S2:**
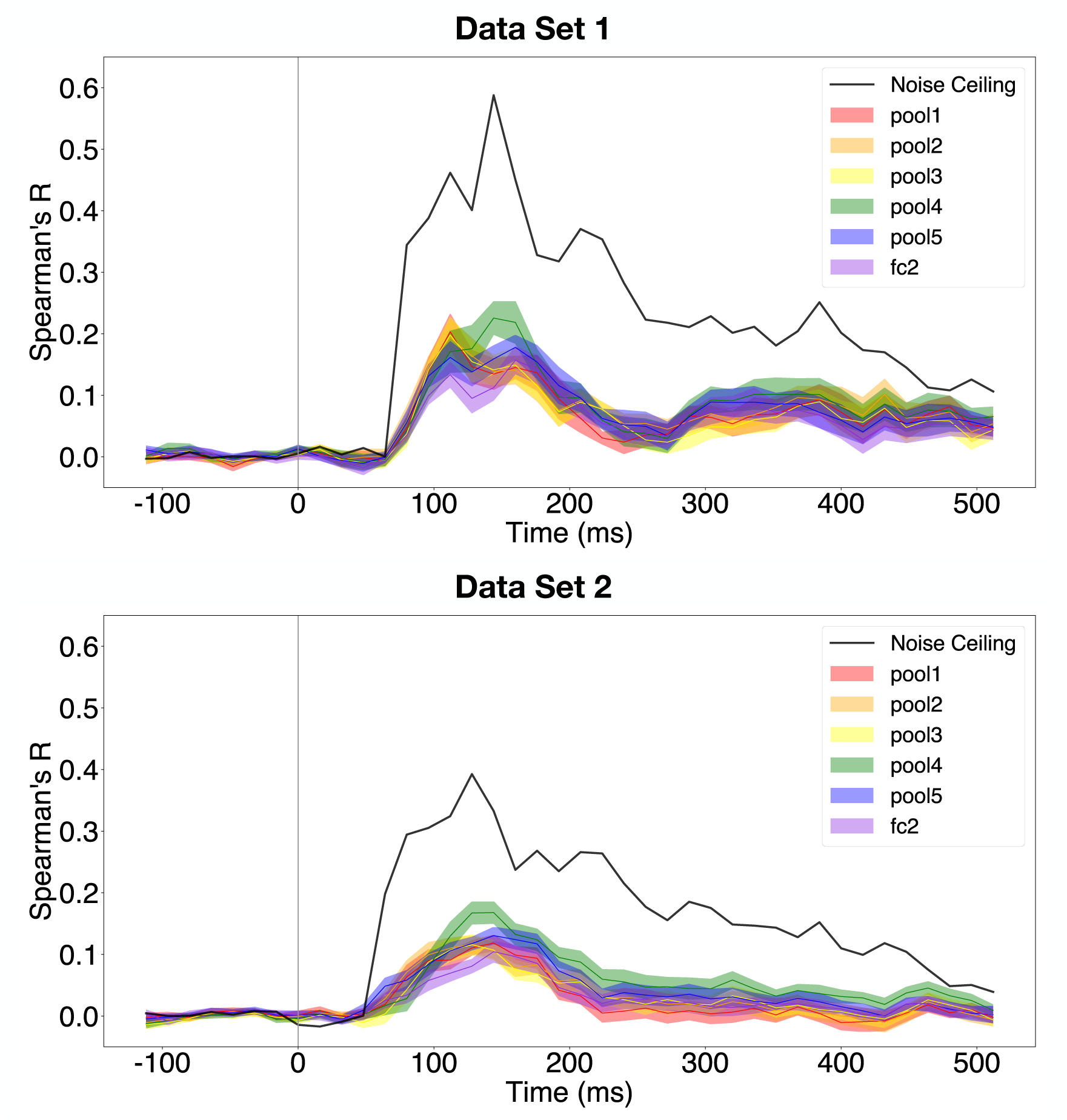
Correlation time courses for all layers using RDMs computed with the cross-validated Pearson correlation metric. Solid lines indicate the mean correlation across subjects and the shaded regions indicate the standard error of the mean across subjects. The vertical grey bar indicates time of stimulus onset. **Top**: Correlation time courses computed using Data Set 1. **Bottom**: Correlation time courses computed using Data Set 2.

**Figure S3:**
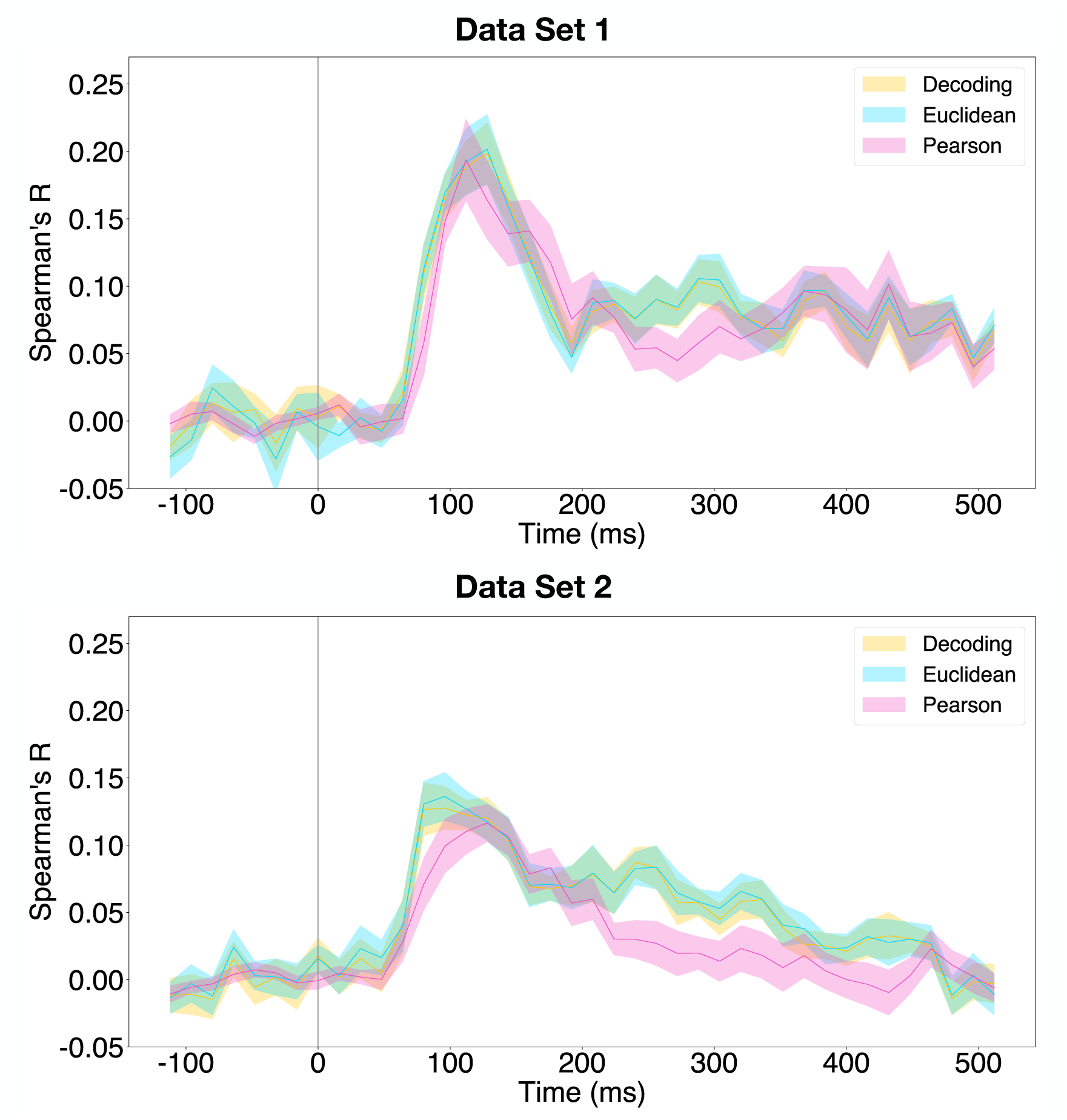
Correlation time course for `pool2` with respect to all three methods. Solid lines indicate the mean correlation across subjects and the shaded regions indicate the standard error of the mean across subjects. The vertical grey bar indicates time of stimulus onset. **Top**: Correlation time courses computed using Data Set 1. **Bottom**: Correlation time courses computed using Data Set 2.

**Figure S4:**
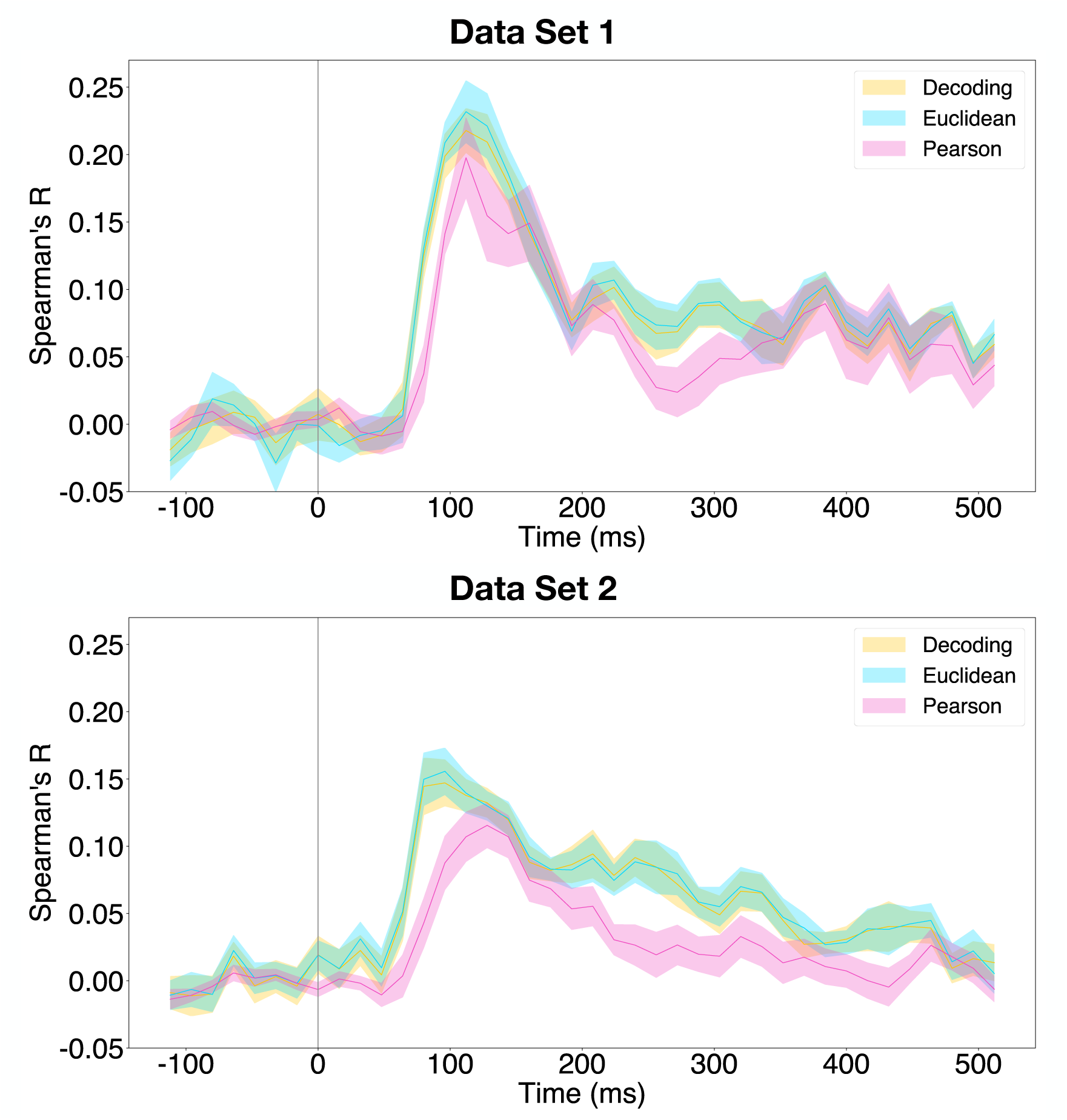
Correlation time course for `pool3` with respect to all three methods. Solid lines indicate the mean correlation across subjects and the shaded regions indicate the standard error of the mean across subjects. The vertical grey bar indicates time of stimulus onset. **Top**: Correlation time courses computed using Data Set 1. **Bottom**: Correlation time courses computed using Data Set 2.

**Figure S5:**
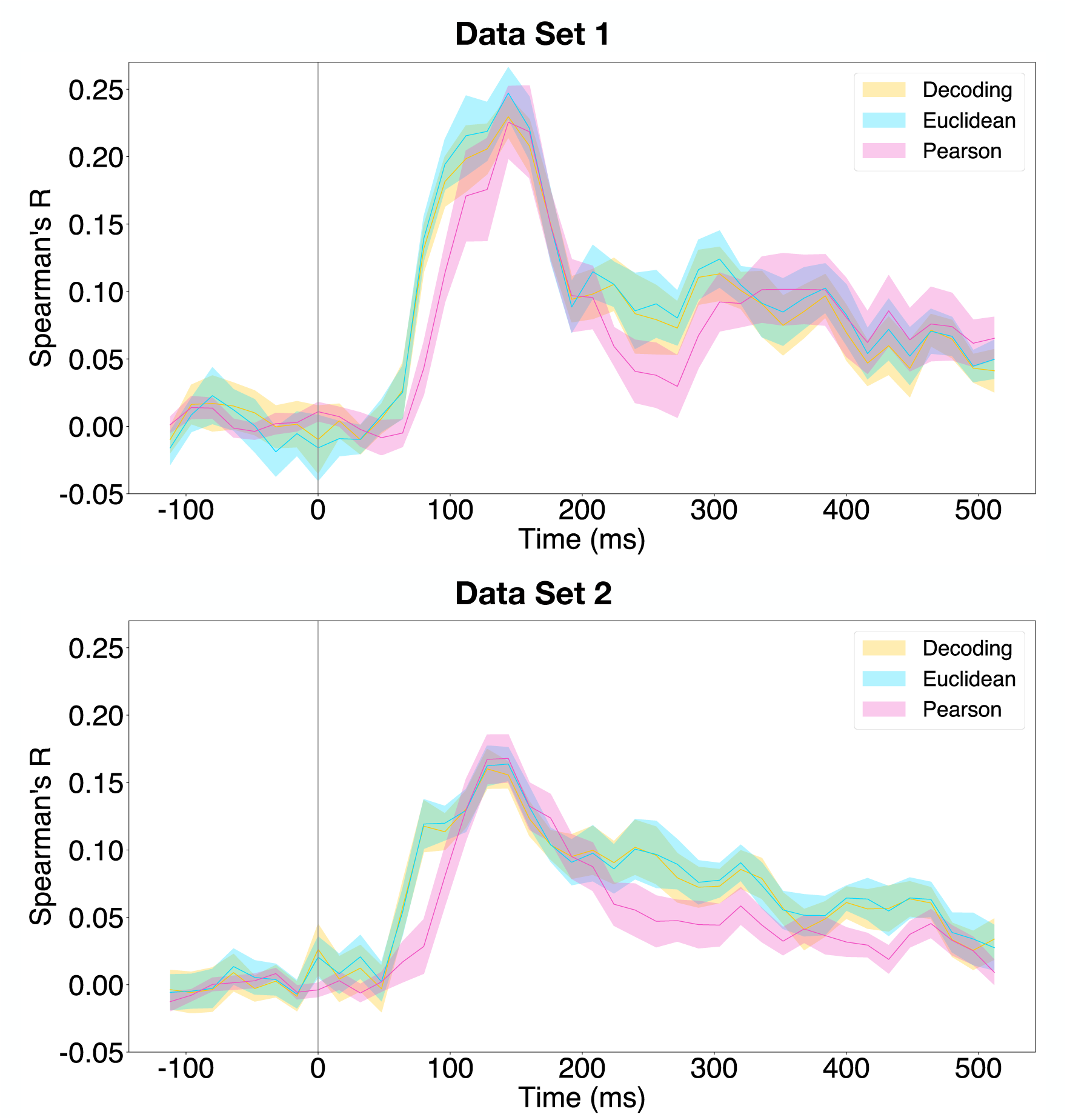
Correlation time course for `pool4` with respect to all three methods. Solid lines indicate the mean correlation across subjects and the shaded regions indicate the standard error of the mean across subjects. The vertical grey bar indicates time of stimulus onset. **Top**: Correlation time courses computed using Data Set 1. **Bottom**: Correlation time courses computed using Data Set 2.

**Figure S6:**
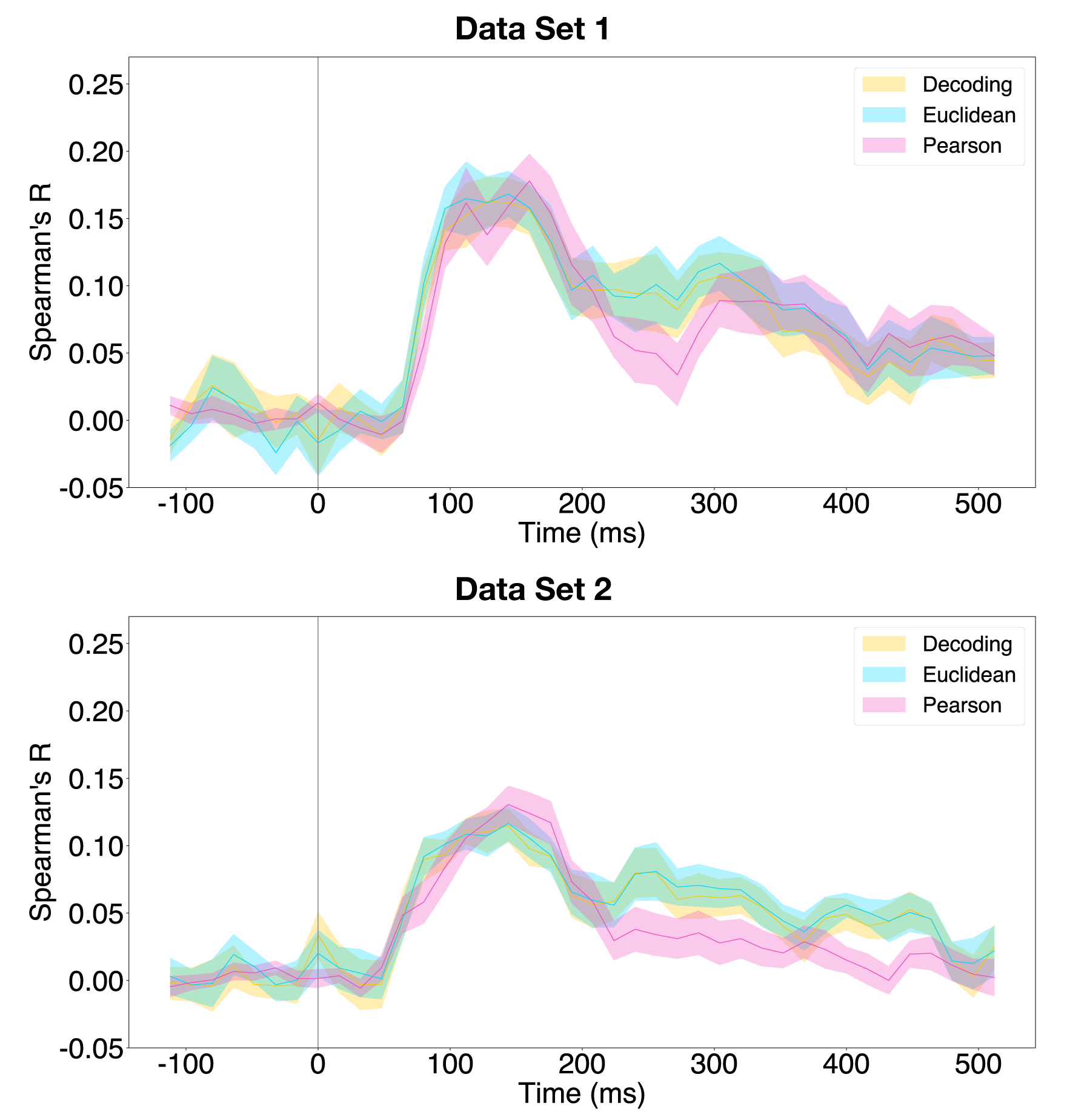
Correlation time course for `pool5` with respect to all three methods. Solid lines indicate the mean correlation across subjects and the shaded regions indicate the standard error of the mean across subjects. The vertical grey bar indicates time of stimulus onset. **Top**: Correlation time courses computed using Data Set 1. **Bottom**: Correlation time courses computed using Data Set 2.

**Figure S7:**
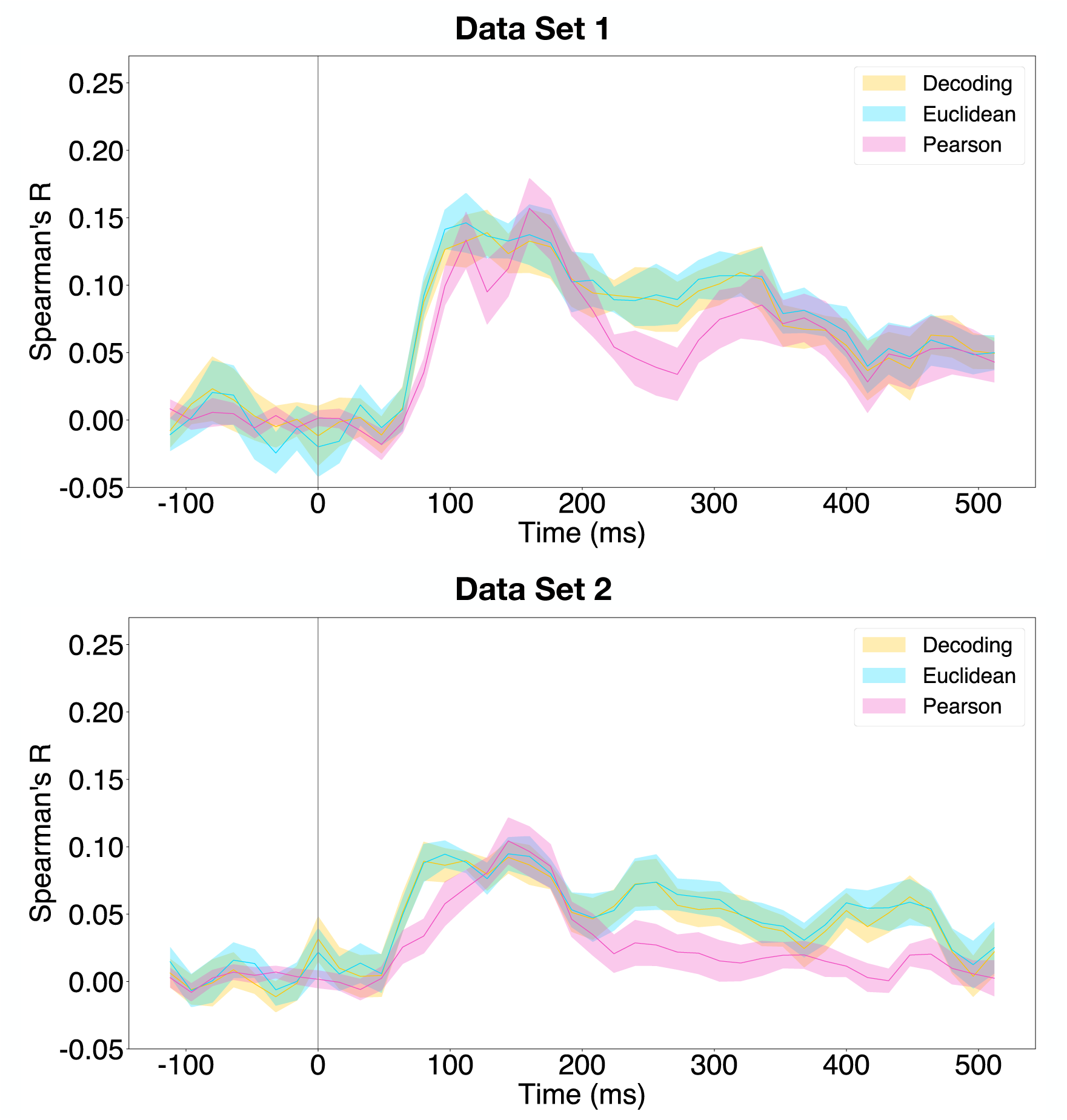
Correlation time course for `fc2` with respect to all three methods. Solid lines indicate the mean correlation across subjects and the shaded regions indicate the standard error of the mean across subjects. The vertical grey bar indicates time of stimulus onset. **Top**: Correlation time courses computed using Data Set 1. **Bottom**: Correlation time courses computed using Data Set 2.

**Figure S8:**
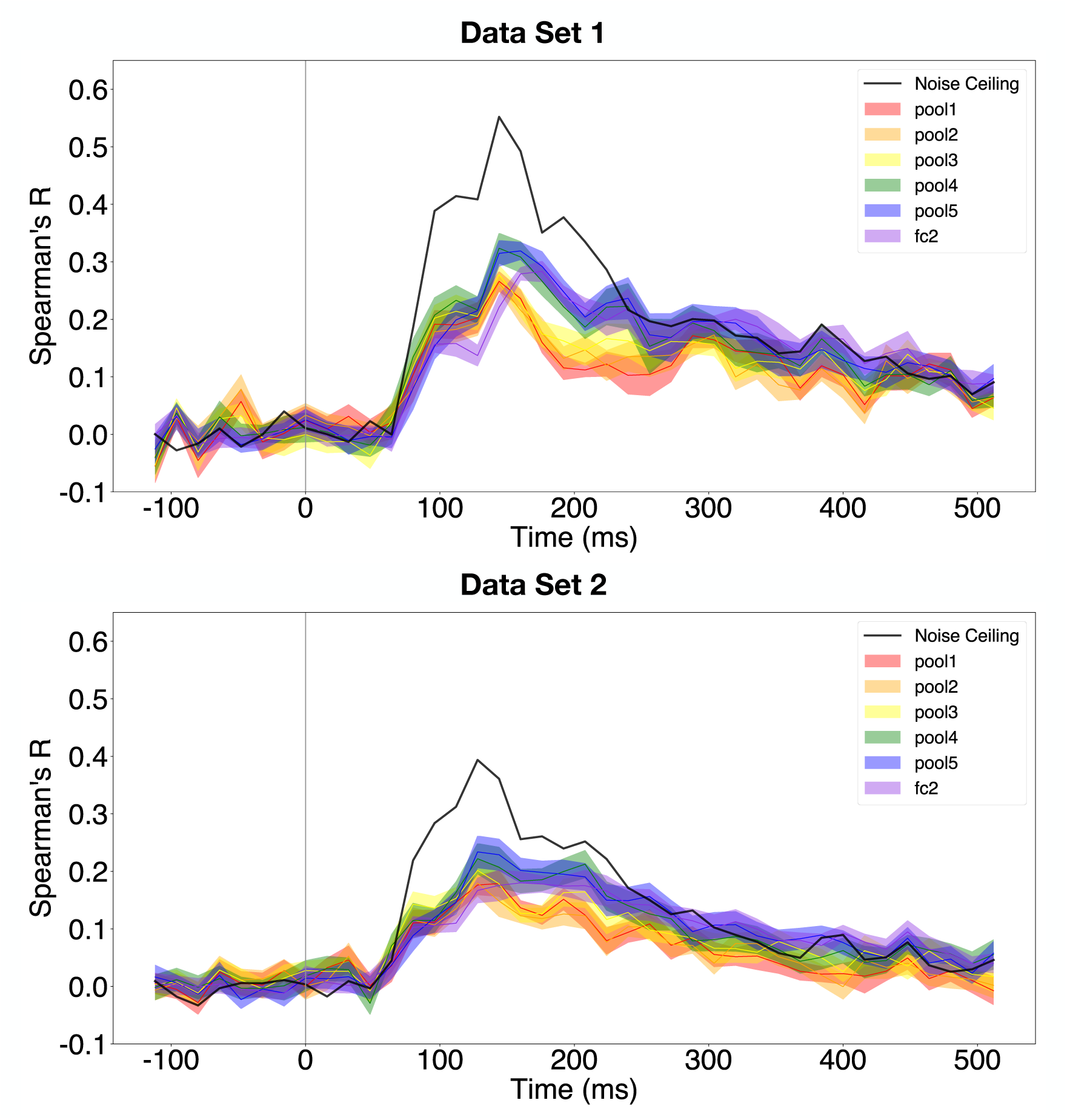
Correlation time courses for all layers using RDMs computed with the decoding accuracy metric and the linear combination optimization procedure. Solid lines indicate the mean correlation across subjects and cross-validation folds and the shaded regions indicate the standard error of the mean across subjects. The vertical grey bar indicates time of stimulus onset. **Top**: Correlation time courses computed using Data Set 1. **Bottom**: Correlation time courses computed using Data Set 2.

**Figure S9:**
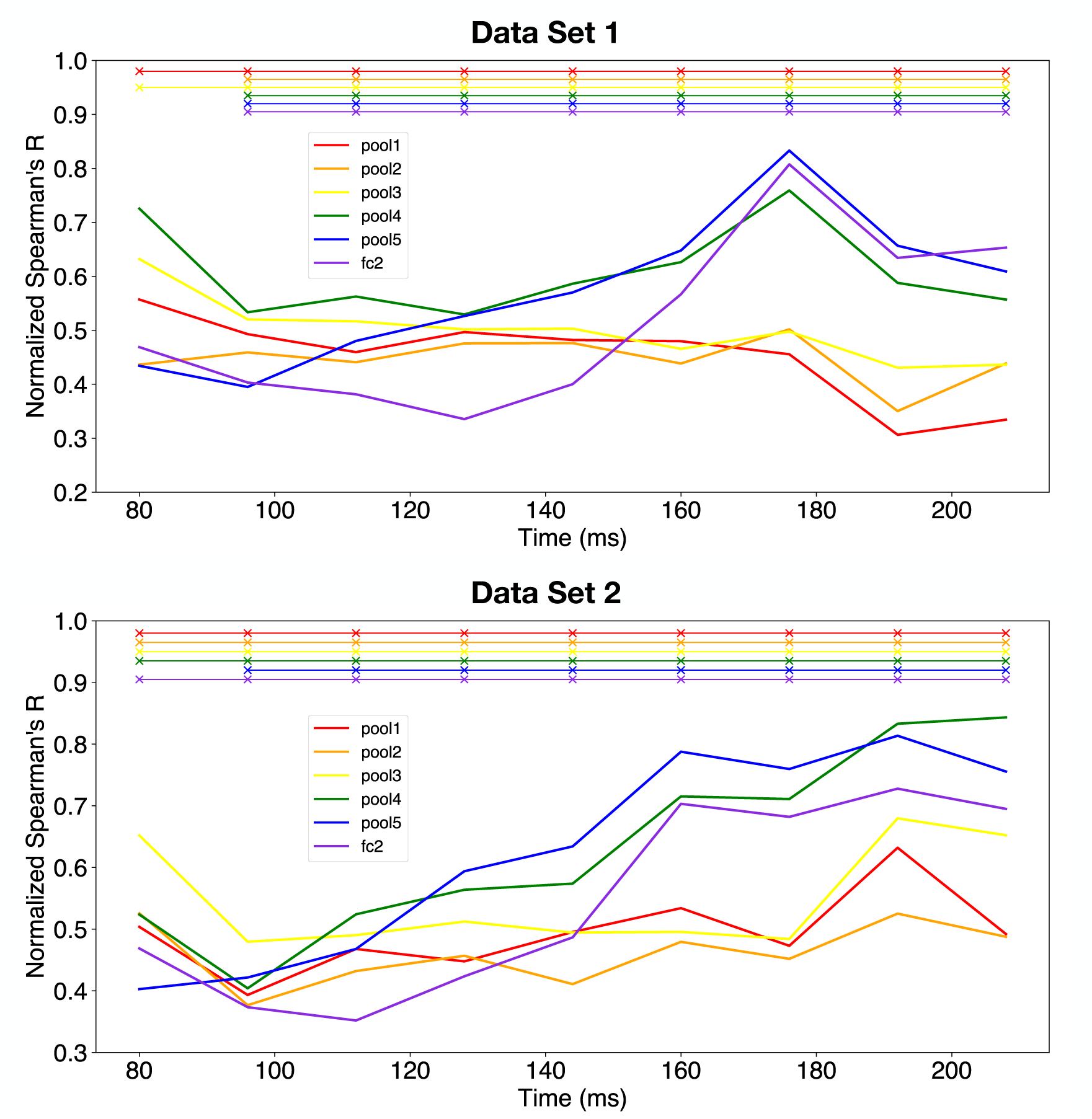
Noise-ceiling normalized correlation time courses for all layers using RDMs computed with the decoding accuracy metric and the linear combination optimization procedure, which were computed by dividing the absolute Spearman’s R by the noise ceiling value at each time point. Solid lines indicate the mean correlation across subjects and cross-validation folds and the *×* symbols at the top of the figure indicate statistically significant correlations (*p <* 0.05, Bonferroni corrected for six comparisons). **Top**: Noise-ceiling normalized correlation time courses computed using Data Set 1. **Bottom**: Noise-ceiling normalized correlation time courses computed using Data Set 2.

**Figure S10:**
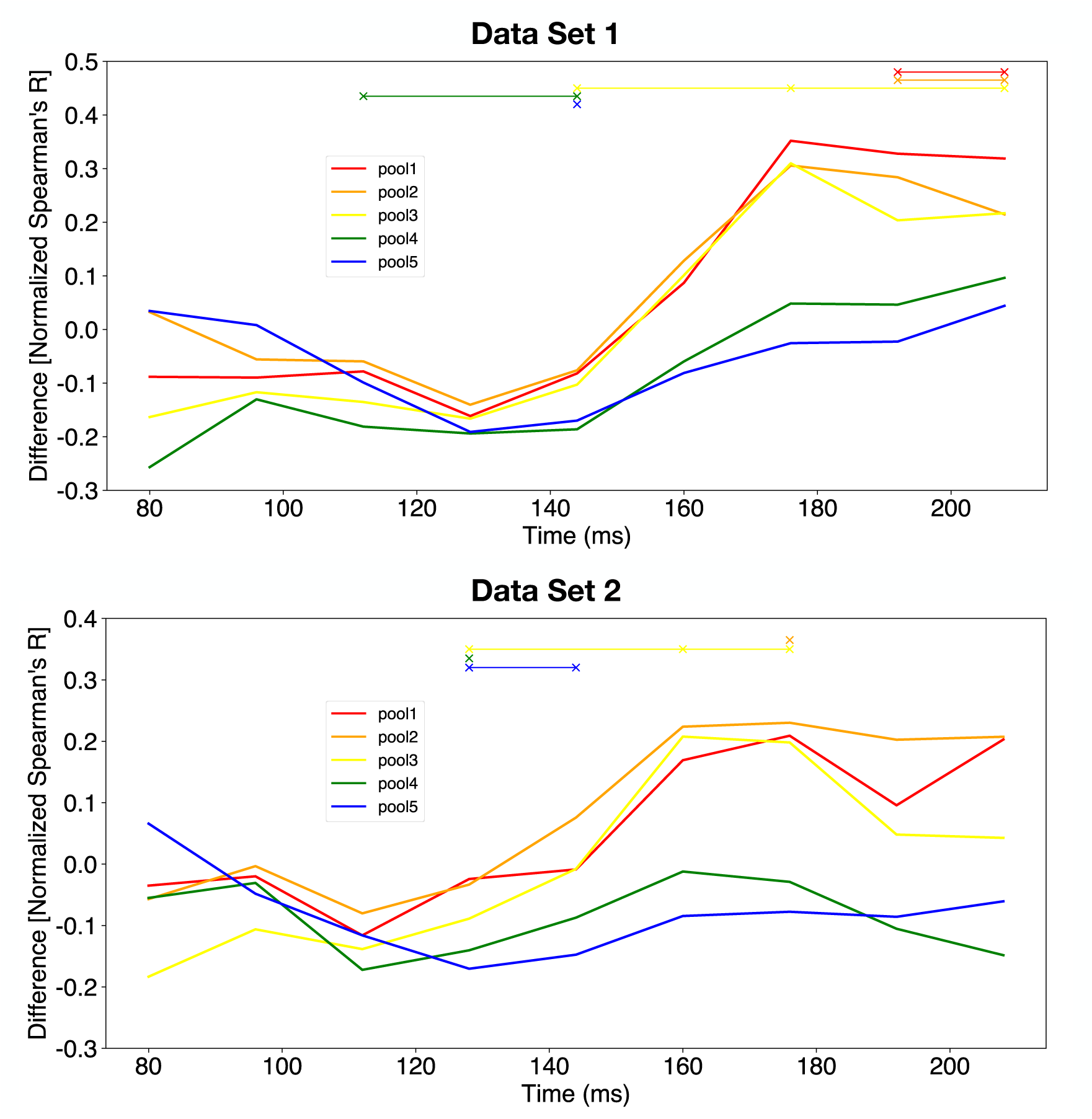
Differences in noise-ceiling normalized correlation time courses for all layers with respect to `fc2` using RDMs computed with the decoding accuracy metric and the linear combination optimization procedure. These were computed by subtracting the noise-ceiling normalized correlations of every other CNN layer from that of `fc2`. Positive values indicate that the correlation value for `fc2` is larger than that of the respective layer. Similarly, negative values indicate that the correlation value for `fc2` is smaller than that of the respective layer. Solid lines indicate the mean correlation across subjects and cross-validation folds and the *×* symbols at the top of the figure indicate statistically significant differences (*p <* 0.05, Bonferroni corrected for five comparisons). **Top**: Differences in noise-ceiling normalized correlation time courses computed using Data Set 1. **Bottom**: Differences in noise-ceiling normalized correlation time courses computed using Data Set 2.

**Figure S11:**
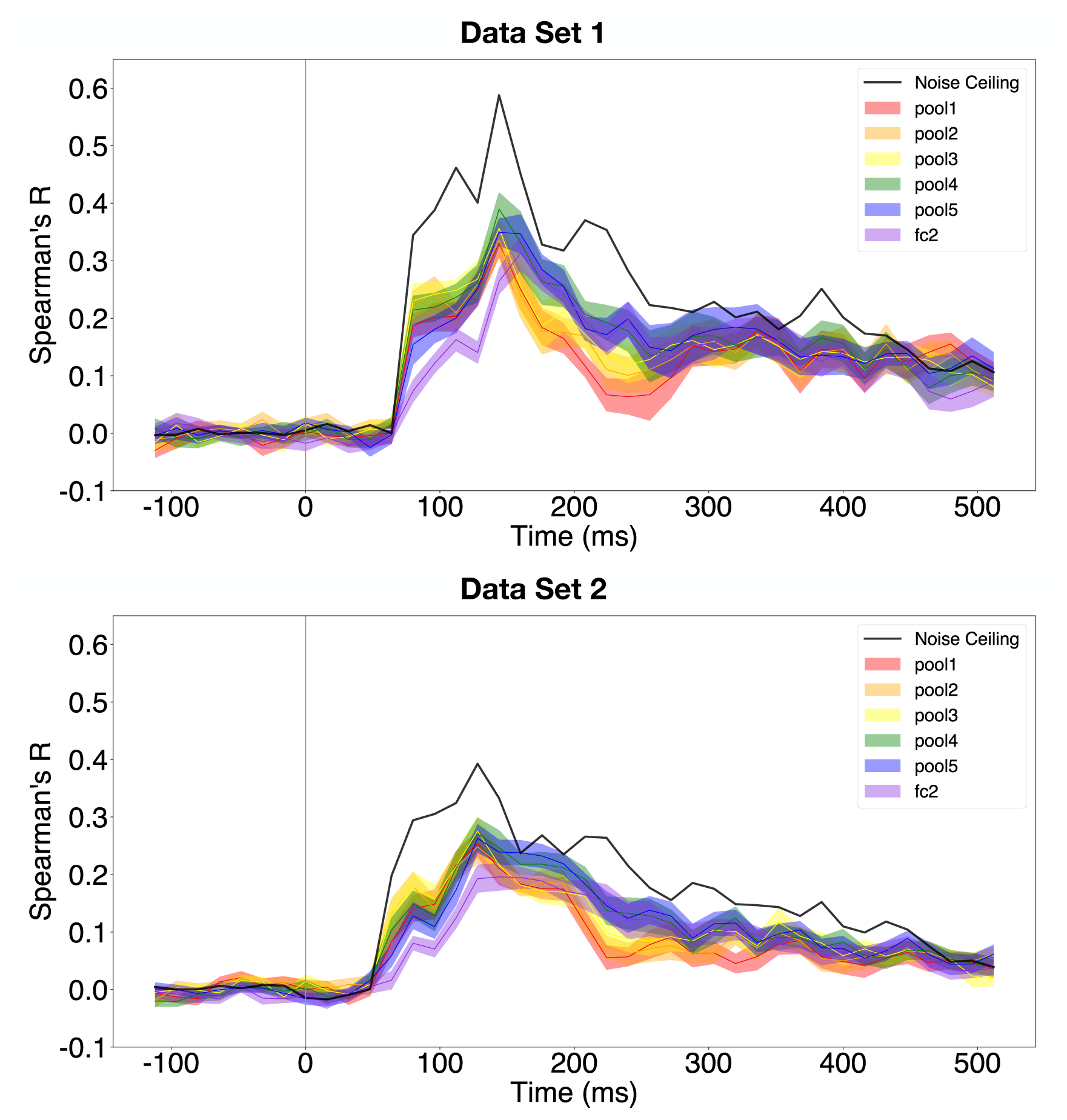
Correlation time courses for all layers using RDMs computed with the cross-validated Pearson correlation metric and the linear combination optimization procedure. Solid lines indicate the mean correlation across subjects and cross-validation folds and the shaded regions indicate the standard error of the mean across subjects. The vertical grey bar indicates time of stimulus onset. **Top**: Correlation time courses computed using Data Set 1. **Bottom**: Correlation time courses computed using Data Set 2.

**Figure S12:**
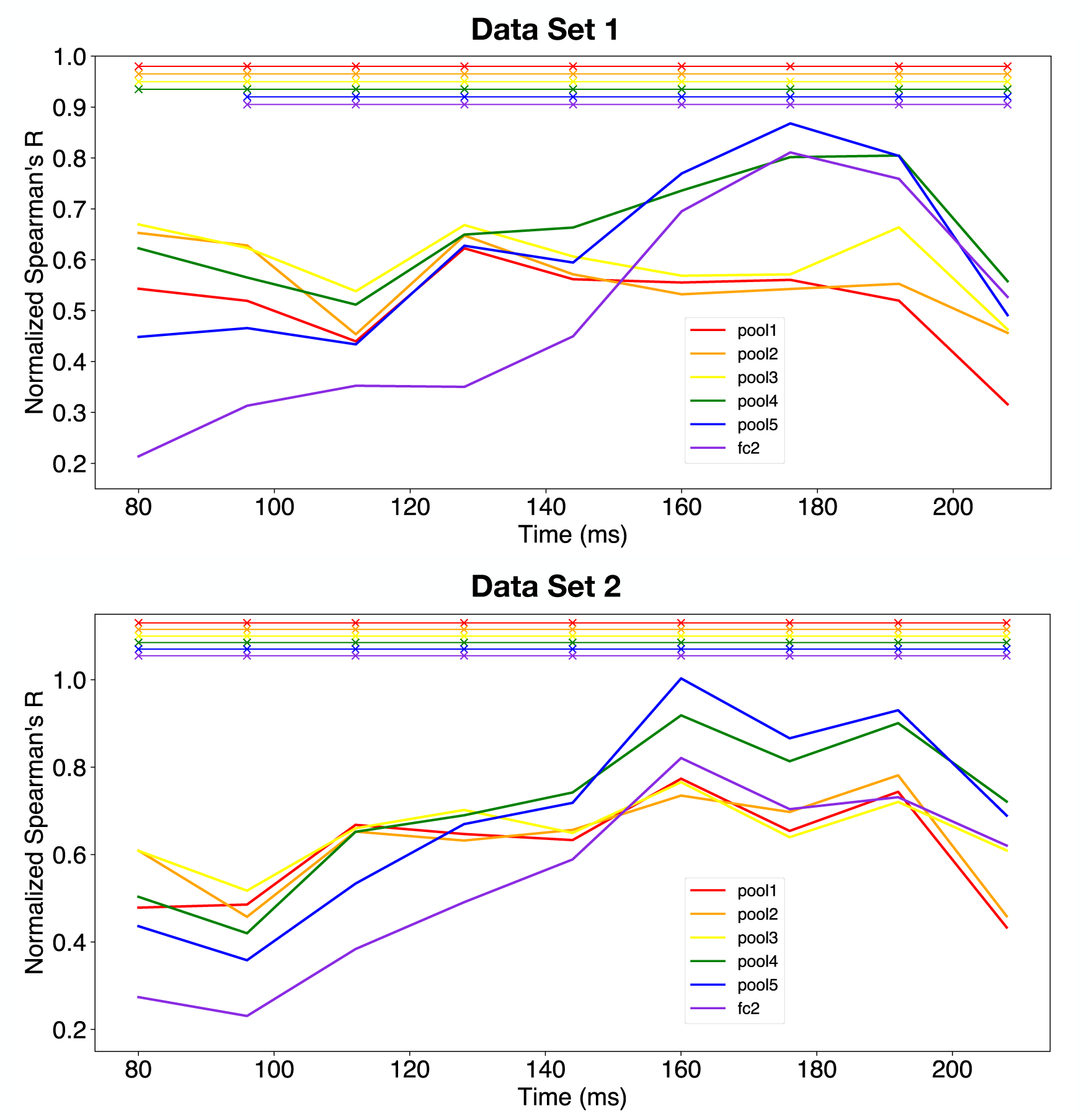
Noise-ceiling normalized correlation time courses for all layers using RDMs computed with the cross-validated Pearson correlation metric and the linear combination optimization procedure, which were computed by dividing the absolute Spearman’s R by the noise ceiling value at each time point. Solid lines indicate the mean correlation across subjects and cross-validation folds and the *×* symbols at the top of the figure indicate statistically significant correlations (*p <* 0.05, Bonferroni corrected for six comparisons). **Top**: Noise-ceiling normalized correlation time courses computed using Data Set 1. **Bottom**: Noise-ceiling normalized correlation time courses computed using Data Set 2.

**Figure S13:**
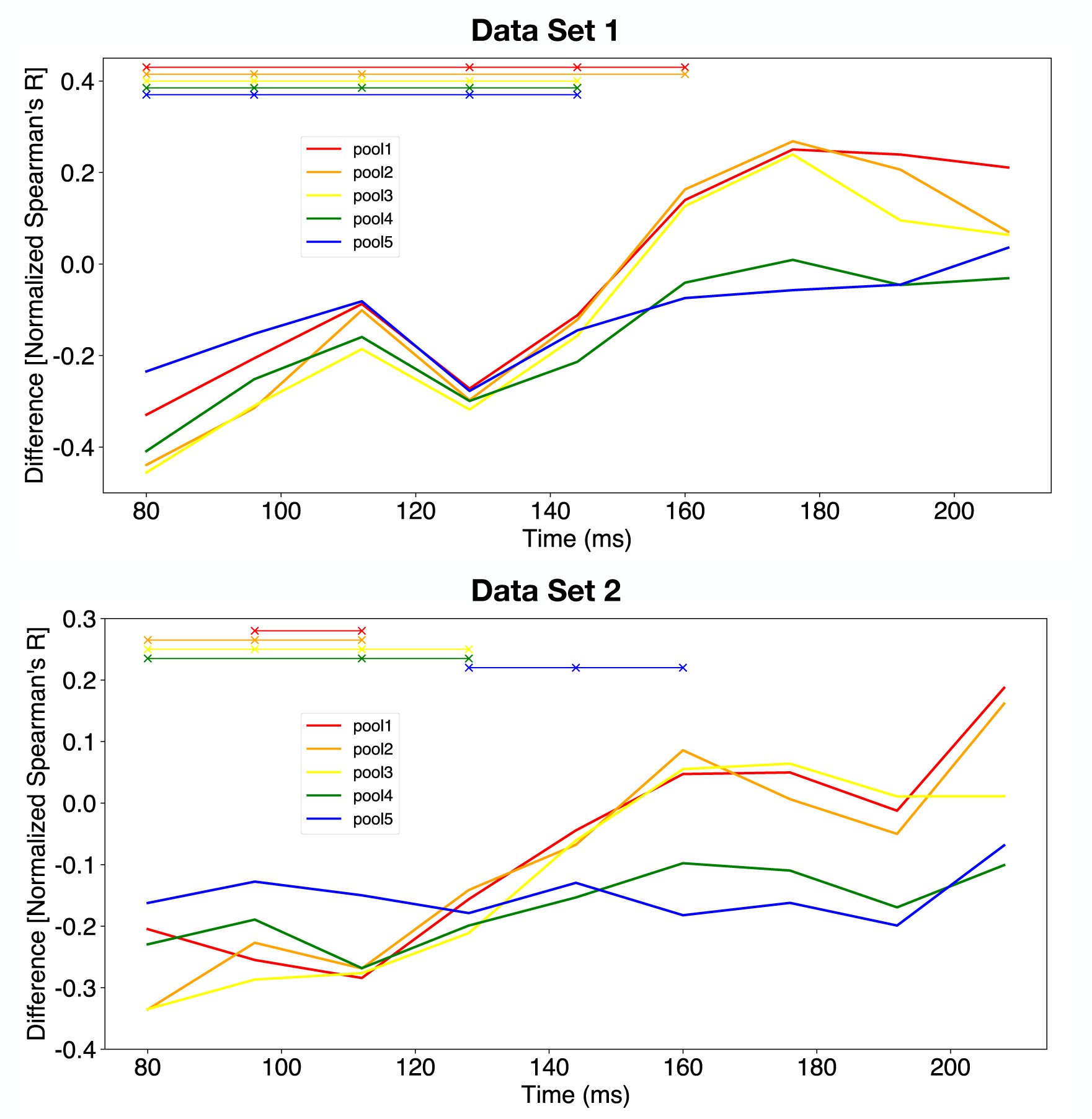
Differences in noise-ceiling normalized correlation time courses for all layers with respect to `fc2` using RDMs computed with the cross-validated Pearson correlation metric and the linear combination optimization procedure. These were computed by subtracting the noise-ceiling normalized correlations of every other CNN layer from that of `fc2`. Positive values indicate that the correlation value for `fc2` is larger than that of the respective layer. Similarly, negative values indicate that the correlation value for `fc2` is smaller than that of the respective layer. Solid lines indicate the mean correlation across subjects and cross-validation folds and the *×* symbols at the top of the figure indicate statistically significant differences (*p <* 0.05, Bonferroni corrected for five comparisons). **Top**: Differences in noise-ceiling normalized correlation time courses computed using Data Set 1. **Bottom**: Differences in noise-ceiling normalized correlation time courses computed using Data Set 2.

**Figure S14:**
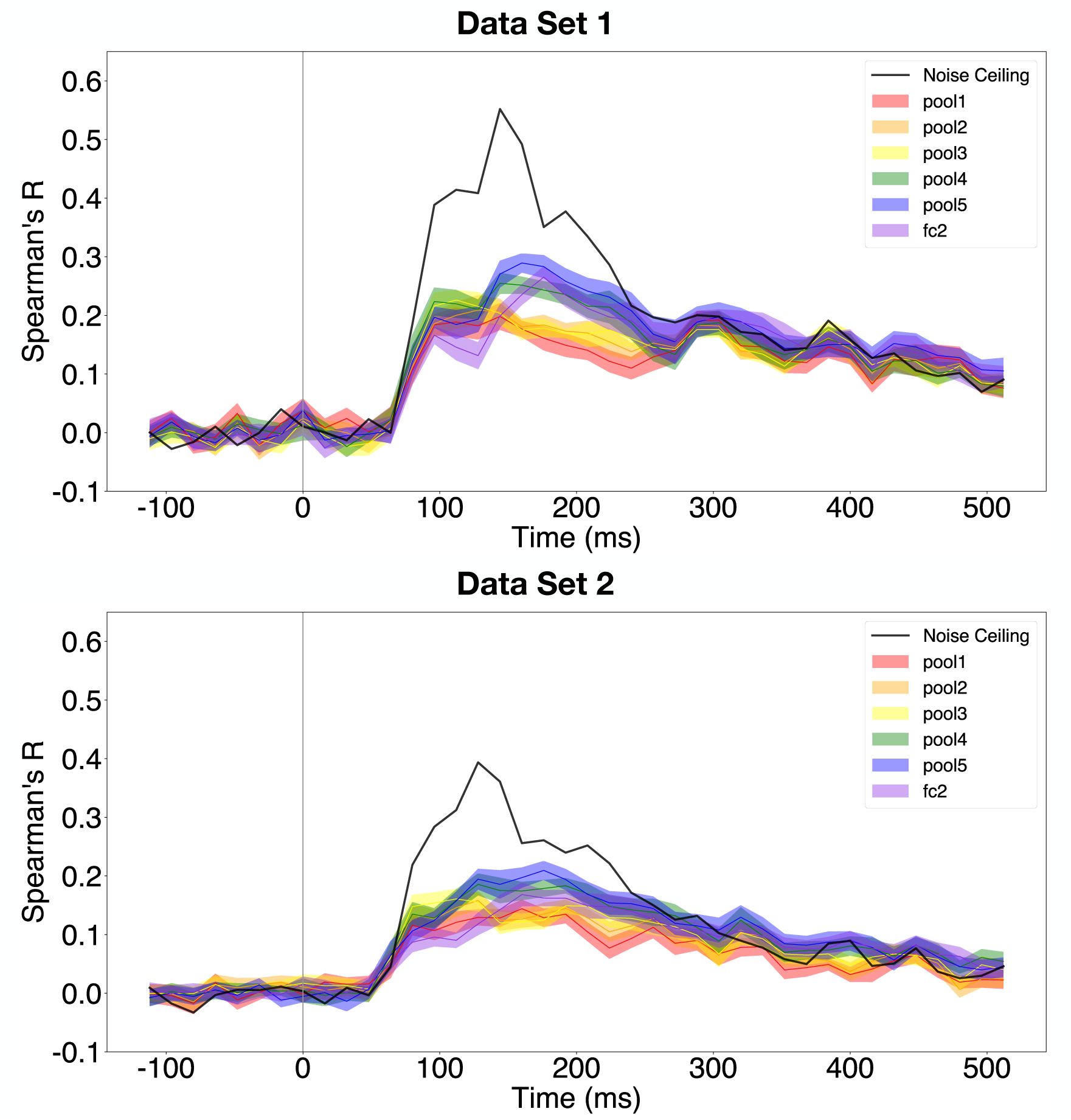
Correlation time courses for all layers using RDMs computed with the decoding accuracy metric and the element-wise scaling optimization procedure. Solid lines indicate the mean correlation across subjects and cross-validation folds and the shaded regions indicate the standard error of the mean across subjects. The vertical grey bar indicates time of stimulus onset. **Top**: Correlation time courses computed using Data Set 1. **Bottom**: Correlation time courses computed using Data Set 2.

**Figure S15:**
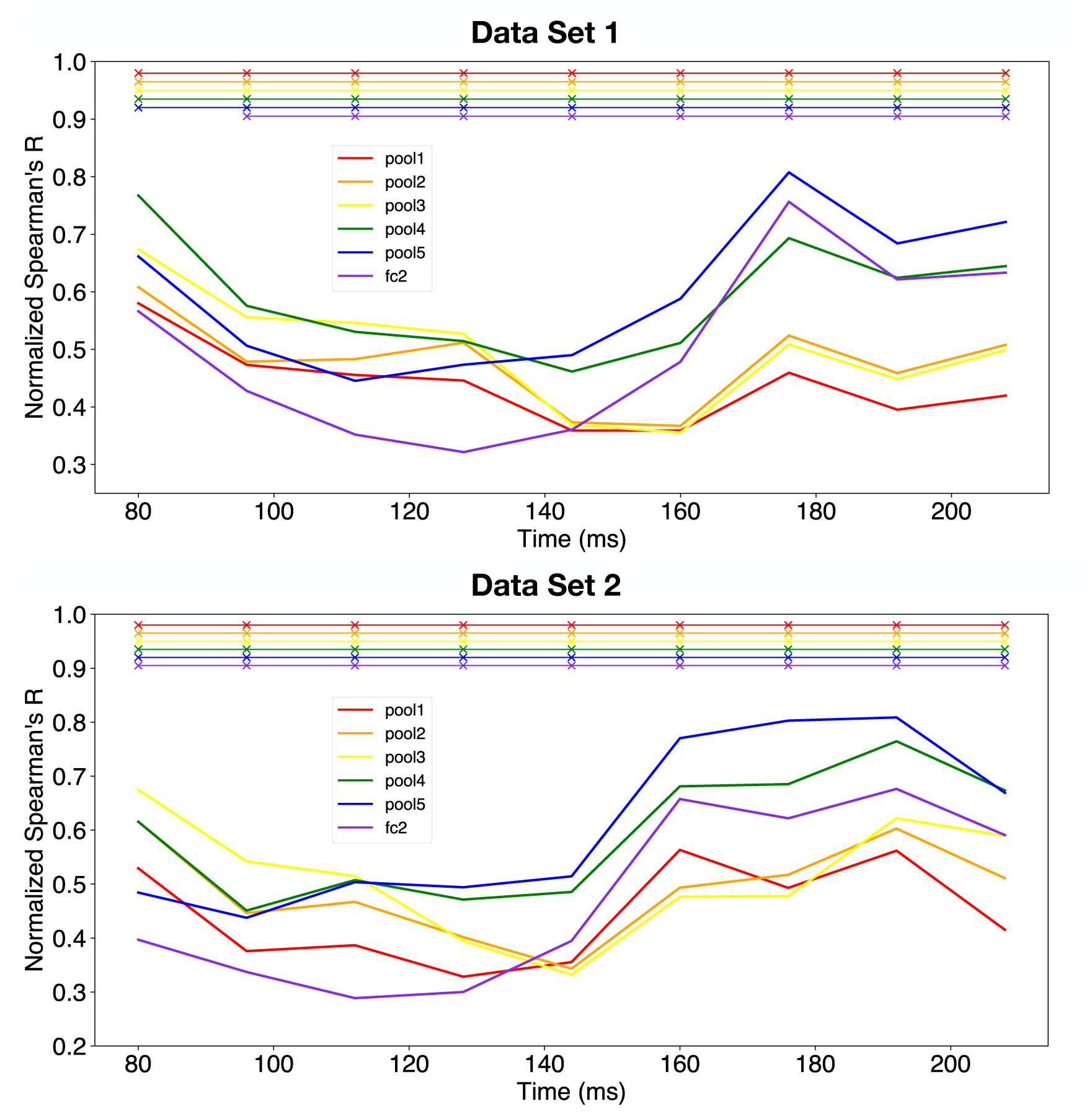
Noise-ceiling normalized correlation time courses for all layers using RDMs computed with the decoding accuracy metric and the element-wise scaling optimization procedure, which were computed by dividing the absolute Spearman’s R by the noise ceiling value at each time point. Solid lines indicate the mean correlation across subjects and cross-validation folds and the *×* symbols at the top of the figure indicate statistically significant correlations (*p <* 0.05, Bonferroni corrected for six comparisons). **Top**: Noise-ceiling normalized correlation time courses computed using Data Set 1. **Bottom**: Noise-ceiling normalized correlation time courses computed using Data Set 2.

**Figure S16:**
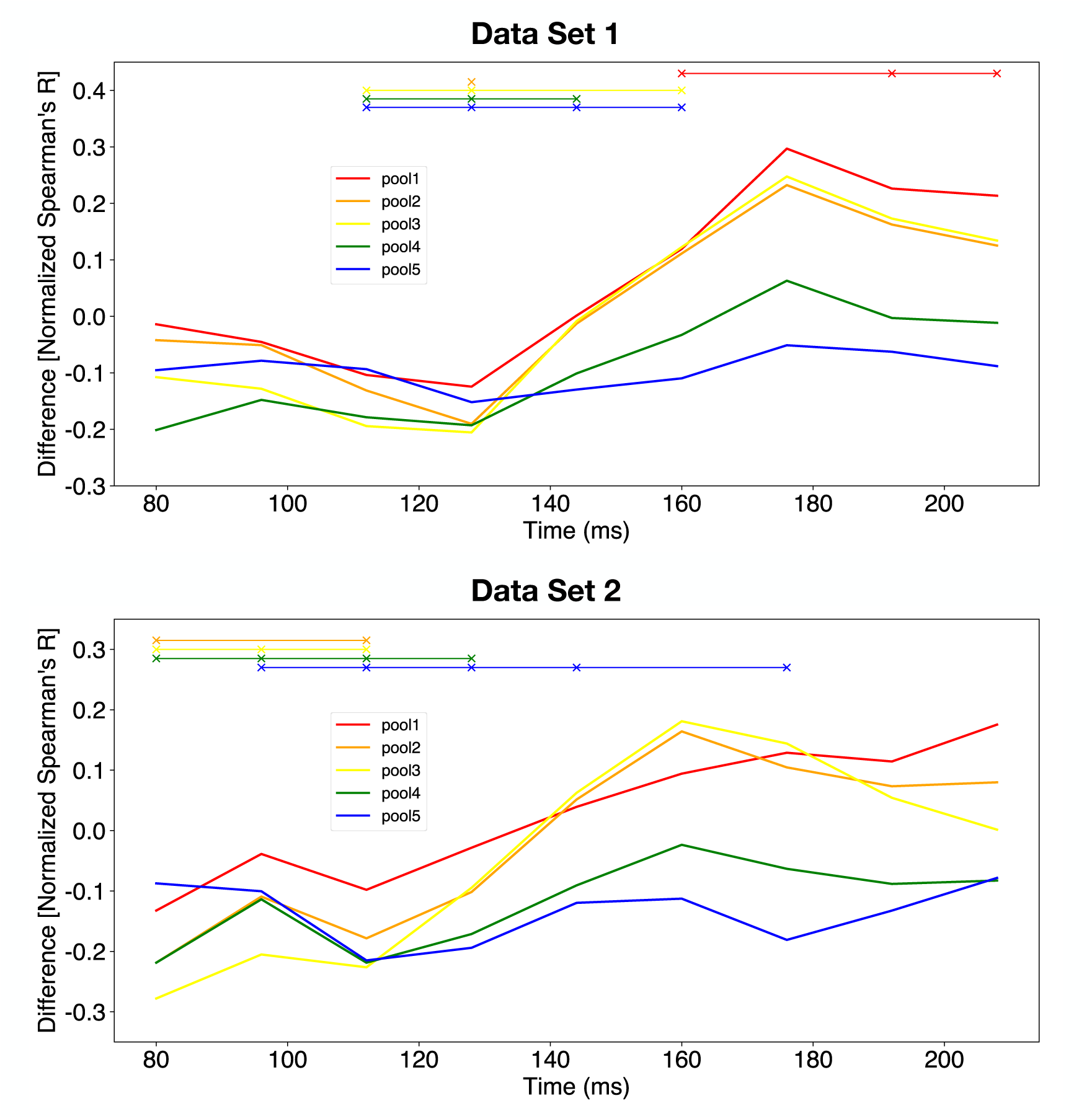
Differences in noise-ceiling normalized correlation time courses for all layers with respect to `fc2` using RDMs computed with the decoding accuracy metric and the element-wise scaling optimization procedure. These were computed by subtracting the noise-ceiling normalized correlations of every other CNN layer from that of `fc2`. Positive values indicate that the correlation value for `fc2` is larger than that of the respective layer. Similarly, negative values indicate that the correlation value for `fc2` is smaller than that of the respective layer. Solid lines indicate the mean correlation across subjects and cross-validation folds and the *×* symbols at the top of the figure indicate statistically significant differences (*p <* 0.05, Bonferroni corrected for five comparisons). **Top**: Differences in noise-ceiling normalized correlation time courses computed using Data Set 1. **Bottom**: Differences in noise-ceiling normalized correlation time courses computed using Data Set 2.

**Figure S17:**
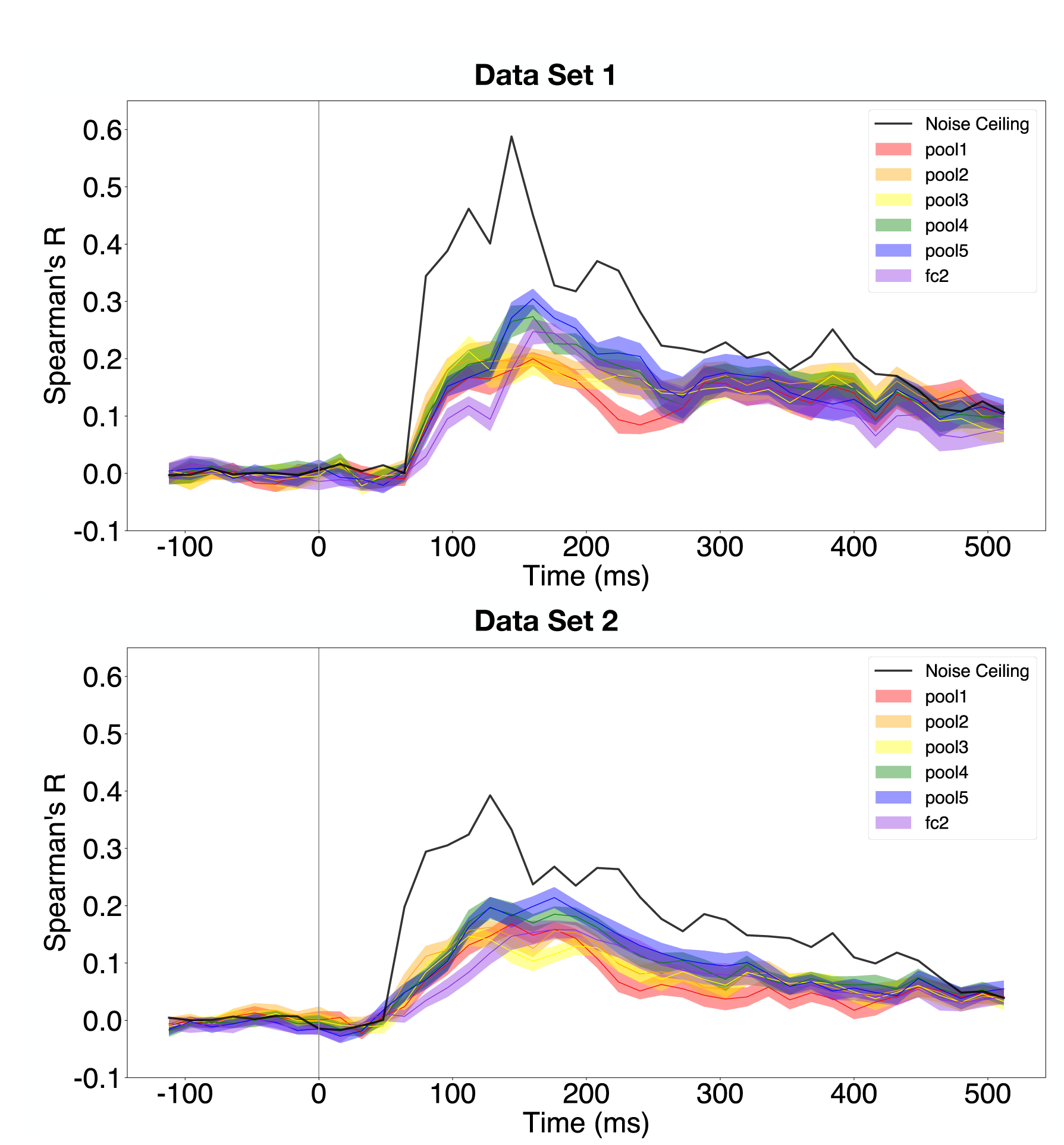
Correlation time courses for all layers using RDMs computed with the cross-validated Pearson correlation metric and the element-wise scaling optimization procedure. Solid lines indicate the mean correlation across subjects and cross-validation folds and the shaded regions indicate the standard error of the mean across subjects. The vertical grey bar indicates time of stimulus onset. **Top**: Correlation time courses computed using Data Set 1. **Bottom**: Correlation time courses computed using Data Set 2.

**Figure S18:**
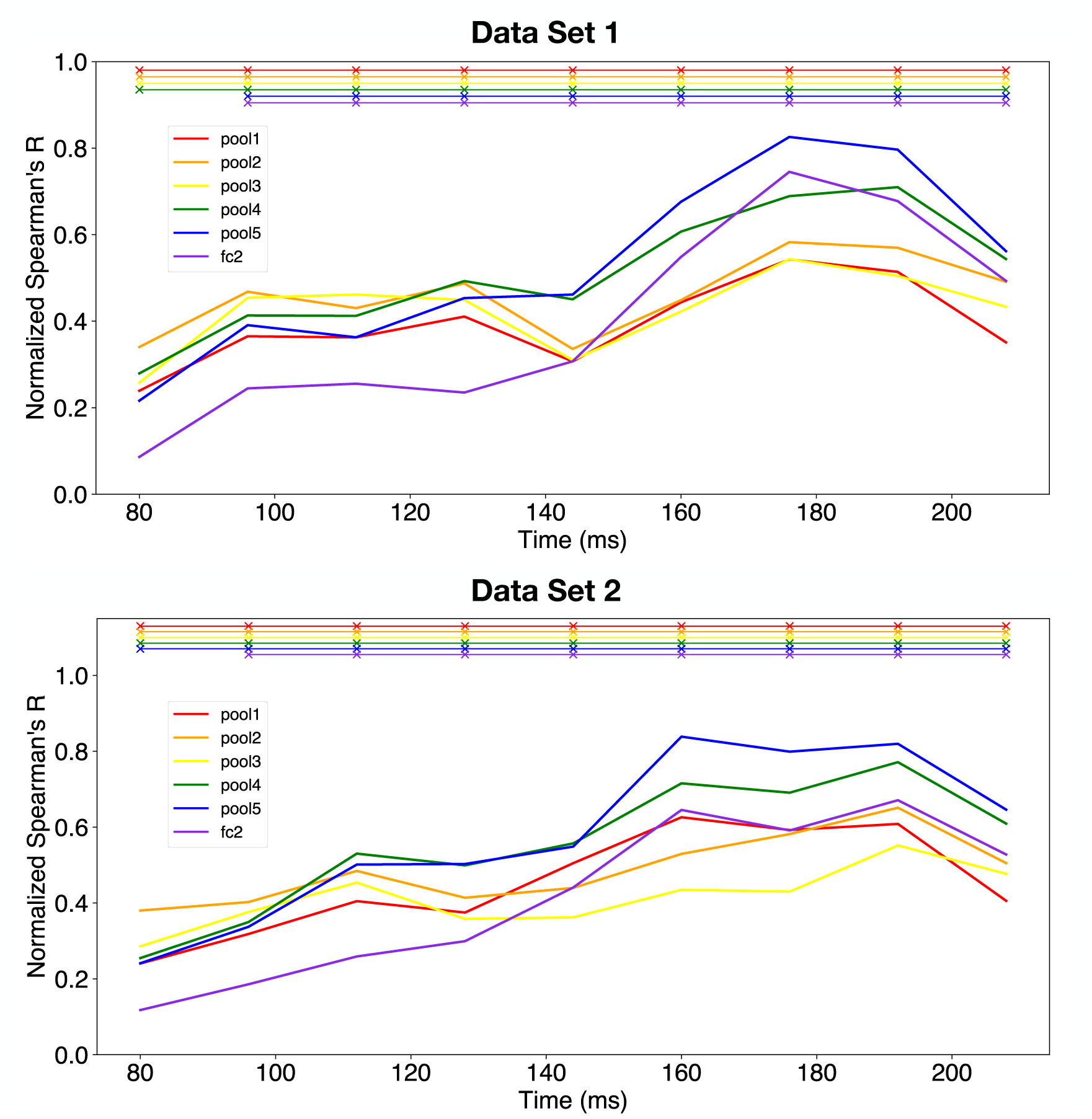
Noise-ceiling normalized correlation time courses for all layers using RDMs computed with the cross-validated Pearson correlation metric and the element-wise scaling optimization procedure, which were computed by dividing the absolute Spearman’s R by the noise ceiling value at each time point. Solid lines indicate the mean correlation across subjects and cross-validation folds and the *×* symbols at the top of the figure indicate statistically significant correlations (*p <* 0.05, Bonferroni corrected for six comparisons). **Top**: Noise-ceiling normalized correlation time courses computed using Data Set 1. **Bottom**: Noise-ceiling normalized correlation time courses computed using Data Set 2.

**Figure S19:**
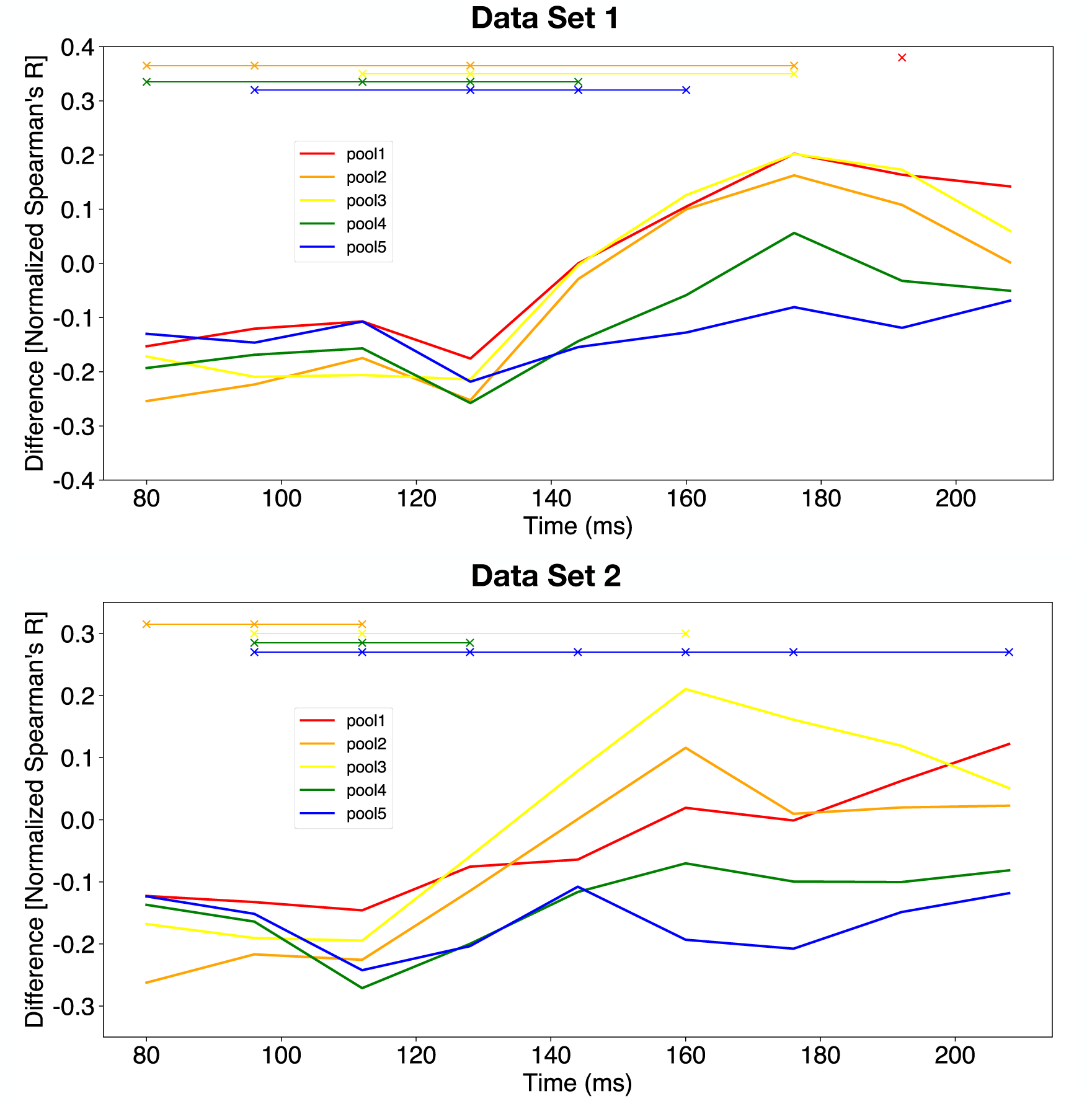
Differences in noise-ceiling normalized correlation time courses for all layers with respect to `fc2` using RDMs computed with the cross-validated Pearson correlation metric and the element-wise scaling optimization procedure. These were computed by subtracting the noise-ceiling normalized correlations of every other CNN layer from that of `fc2`. Positive values indicate that the correlation value for `fc2` is larger than that of the respective layer. Similarly, negative values indicate that the correlation value for `fc2` is smaller than that of the respective layer. Solid lines indicate the mean correlation across subjects and cross-validation folds and the *×* symbols at the top of the figure indicate statistically significant differences (*p <* 0.05, Bonferroni corrected for five comparisons). **Top**: Differences in noise-ceiling normalized correlation time courses computed using Data Set 1. **Bottom**: Differences in noise-ceiling normalized correlation time courses computed using Data Set 2.

